# Polo-like kinase-1 Inhibitors and the Antiandrogen Abiraterone Synergistically Disrupt Mitosis and Kill Cancer Cells of Disparate Origin Independently of Androgen Receptor Signaling

**DOI:** 10.1101/2022.05.12.491633

**Authors:** Jesse C. Patterson, Andreas Varkaris, Peter J. P. Croucher, Maya Ridinger, Susan Dalrymple, Mannan Nouri, Fang Xie, Shohreh Varmeh, Oliver Jonas, Matthew A. Whitman, Sen Chen, Saleh Rashed, Lovemore Makusha, Jun Luo, John T. Isaacs, Mark G. Erlander, David J. Einstein, Steven P. Balk, Michael B. Yaffe

## Abstract

Abiraterone, a standard treatment for metastatic castrate-resistant prostate cancer (mCRPC), slows disease progression by abrogating androgen synthesis and antagonizing the androgen receptor (AR). We report that inhibitors of the mitotic kinase Plk1, including the clinically active third-generation Plk1 inhibitor onvansertib, when co-administered with abiraterone, synergistically kill cancer cells from a wide variety of tumor types in an androgen-independent manner, both *in vitro* and *in vivo*. Abiraterone treatment alone results in defects in mitotic spindle orientation, failure of complete chromosome condensation, and upregulation of mitosis and mitotic-spindle related gene sets independently of its effects on AR signaling. These effects, while mild following abiraterone monotherapy, result in profound sensitization to the anti-mitotic effects of Plk1 inhibition, leading to spindle assembly checkpoint-dependent mitotic cell death and entosis. In a murine PDX model of mCRPC, combined onvansertib and abiraterone resulted in enhanced mitotic arrest and dramatic inhibition of tumor cell growth compared to either agent alone.

**STATEMENT OF SIGNIFICANCE:** A phase 2 clinical trial is underway (NCT03414034) testing combined Plk1 inhibitor onvansertib and abiraterone in mCRPC patients with nascent abiraterone resistance. Our work establishes a mechanistic basis for that trial and indicates that combined abiraterone and onvansertib co-treatment may have broad utility for cancer treatment beyond mCRPC.

## INTRODUCTION

Prostate cancer is largely driven by signaling through the androgen receptor (AR), a nuclear receptor that binds both its ligand testosterone and chromatin to directly regulate gene transcription. Prior to androgen stimulation, the AR is located in the cytoplasm, while upon ligand binding it translocates to the nucleus, where it binds to AR-response elements on a subset of genes that drive tumor cell proliferation and upregulate survival pathways (1). Accordingly, first-line systemic therapy for prostate cancer is administered to men for whom surgical resection or radiotherapy is insufficient for tumor control, and targets signaling through the AR by suppressing the endogenous production of androgens within the testes (androgen deprivation therapy; ADT). While ADT is initially effective for the vast majority of men, patients invariably develop metastatic castration-resistant prostate cancer (mCRPC). This occurs through multiple mechanisms including amplification of the AR gene, expression of constitutively active AR splice variants (AR-v7), increased crosstalk between the AR and other signaling pathways, and upregulation of intratumoral androgen synthesis (2). Treatment of CPRC involves the use of the antiandrogens abiraterone acetate or enzalutamide to further suppress AR signaling. Abiraterone, a synthetic derivative of the endogenous steroid pregnenolone, abrogates both residual adrenal and intratumoral androgen synthesis by inhibiting Cyp17A1 (**Supplemental Figure 1A**) (3, 4). In addition, both abiraterone and its metabolic derivative Δ-4-abiraterone also directly inhibit the AR by interacting with its ligand binding domain (5, 6). Enzalutamide is a direct AR antagonist that inhibits AR signaling irrespective of androgen synthesis (7). Treatment of CRPC patients with either abiraterone or enzalutamide transiently arrests further tumor progression; however, most mCRPC patients develop acquired resistance to these antiandrogens with a median progression-free survival of 1-2 years (8, 9).

Several other signaling pathways cross-talk with the AR pathway, including the PI3-kinase/Akt pathway, the Erk MAP kinase pathway, and WNT/*β*-catenin signaling, among others, to modulate tumor cell survival, invasion, and metastasis (10), as well as confer network robustness that contributes to prostate cancer development (11–13) and resistance to anti-cancer therapies (14, 15). In addition, two other signaling pathways that have been shown to contribute to prostate cancer progression and therapeutic resistance are the Polo-like kinase 1 (Plk1) mitotic kinase pathway (16, 17), and components of the DNA damage response (DDR) signaling pathway (18). Multiple components of the DDR pathway are mutated in prostate cancer including *BRCA1, BRCA2*, MSH2, MSH6 PMS2 and *NBS1*, have been shown to be mutated in prostate cancer or germline mutations increase incidence encoding proteins involved in homologous recombination, as well as several genes encoding proteins involved in mismatch repair (MSH2, MSH6, MLH1, and PMS2) have been reported to increase the risk of prostate cancer development, and has been associated with higher grade tumors and a more aggressive clinical course (19–23). In addition, germline or somatic prostate tumor mutations in other proteins involved in the DNA damage response (RAD51, ATM, CHEK2, PALB2) have been shown to result in non-indolent and lethal disease (24–29).

Plk1 is a serine/threonine kinase that plays a critical role in nearly every stage of mitosis (30, 31). While Plk1 is undetectable in most differentiated non-proliferating adult tissues, it is overexpressed and associated with poor prognosis in many cancers including prostate cancer, in which it is overexpressed in >50% of patients and associated with a higher tumor grade (32, 33). Plk1 is thought to be particularly important for mitotic cell division in cancer cells due to elevated levels of replicative stress and underlying chromosomal instability (34, 35). Overexpression of Plk1 in prostate epithelial cells has been shown to result in malignant transformation, enhanced cell migration and an epithelial to mesenchymal transition mediated, in part, through activation of Erk (36). In addition, Plk1 has been shown to phosphorylate and suppress the pro-apoptotic function of the FOXO1 transcription factor in advanced prostate cancer cells (37) and postulated to cross-talk with the Wnt/β-catenin pathway in CRPC (38).

Synergistic combination therapies are of particular interest due to their potential for enhancing efficacy and cancer cell selectivity, overcoming drug resistance, and possibly lowering toxicity though decreased dosage of individual drugs (39). Given the emerging importance of the PLK1 and DDR signaling pathways in prostate cancer, we sought to investigate whether the introduction of anti-mitotic drugs, or the induction of genotoxic stress, in combination with inhibition of signaling through the androgen receptor, could be used to synergistically enhance anti-tumor responses in ADT-resistant mCRPC cells. We report here the surprising discovery that synergistic tumor-cell killing by combinations of Plk1inhibitors and abiraterone was completely independent of androgen receptor signaling, could be observed in a variety of cancer cell types extending far beyond prostate cancer, and was not recapitulated by combining abiraterone with inhibitors of other mitotic kinases except Plk1. We show that abiraterone treatment alone causes defects in chromosome condensation and mitotic spindle orientation, which results in synergistic mitotic cancer cell death and entosis when abiraterone is combined with inhibitors of Plk1.

## RESULTS

### The antiandrogen abiraterone acetate in combination with inhibitors of Polo-like kinase 1 synergistically kills CRPC cells

To explore pathways in CRPC cells that could be targeted to create abiraterone-based synergistic drug combinations, we performed a combinatorial dose-response study with abiraterone focused on anti-mitotic agents and DNA damaging drugs that could be rapidly translated into the clinic. Docetaxel was investigated because it is an alternative first-line treatment for mCRPC, and is often used as second-line agent after the development of antiandrogen resistance (40). Doxorubicin was examined because it has been shown to synergize with docetaxel in prostate cancer (41). Furthermore, Polkinghorn et al. reported that the AR regulates DNA repair pathways, to promote resistance to DNA damage (42). An inhibitor of the mitotic kinase Plk1 (BI2536) was explored because of our lab’s long-standing interest in how this kinase controls mitotic progression and silences the G2/M checkpoint after DNA damage (43–45), particularly by being targeted to its substrates through phospho-priming and Polo-box dependent interactions (46–48). There are also reports suggesting that during development of CRPC the AR selectively activates genes involved in mitosis, that CRPC cells are particularly sensitive to Plk1 inhibitors, and that AR splice variants that are associated with antiandrogen resistance specifically upregulate genes involved in mitosis including Plk1 (16,49,50). Based on our previous work indicating that the timing of combination drug addition may be a critical parameter in cancer cell phenotypic responses (51), we examined drug combination synergy both when abiraterone was added simultaneously with a second drug, as well as when drugs were administered in a time staggered manner (**Figure 1A**). To mimic levels of androgen observed following castration, C4-2 CRPC cells were grown in charcoal-stripped fetal bovine serum (csFBS) and subjected to combinatorial dose matrices consisting of abiraterone and the drugs listed above.

**Figure 1.**
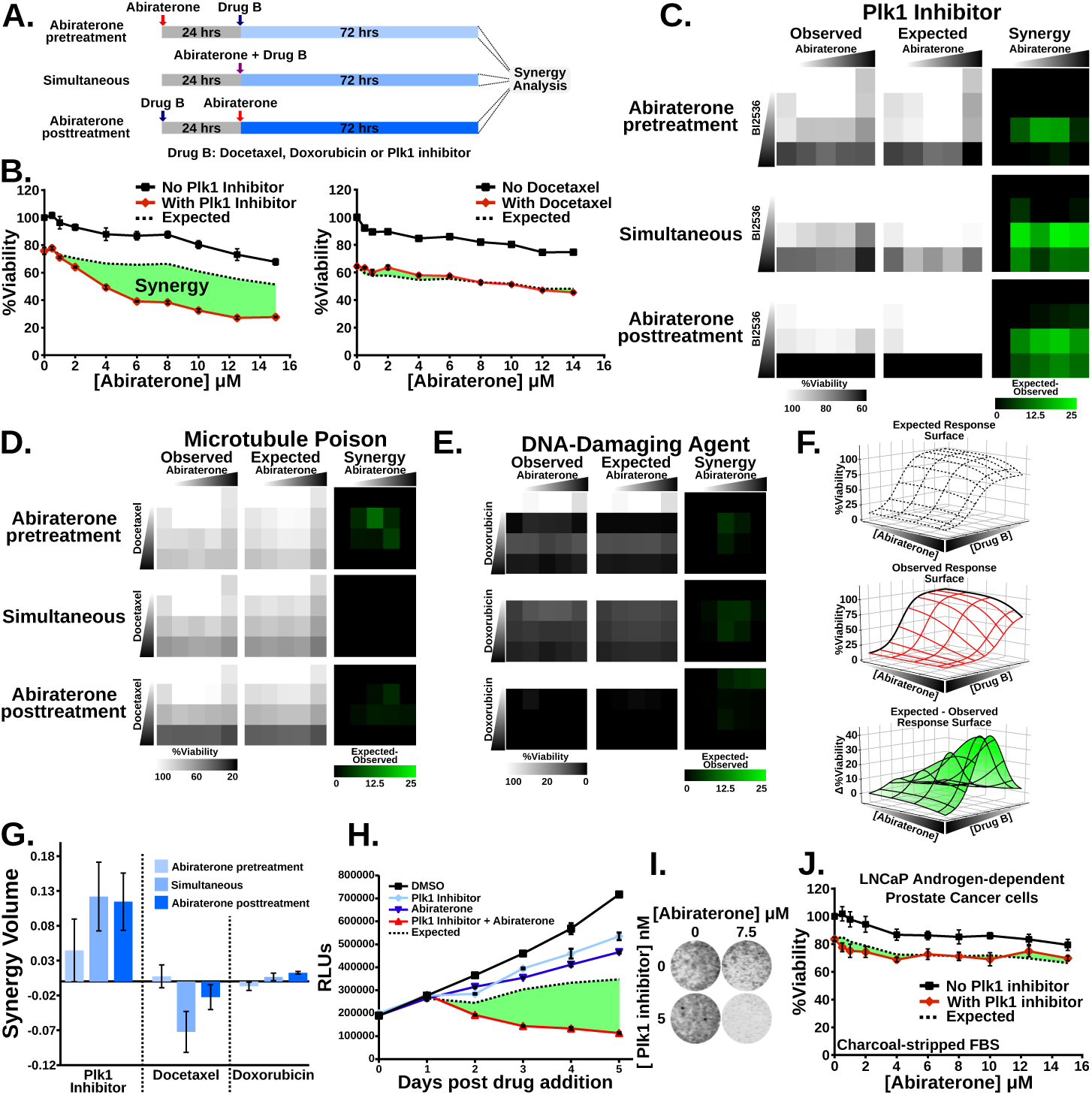
The combination of Plk1 inhibitors and abiraterone synergistically kill CRPC cells. **(A)** Schematic indicating drug testing for synergy with abiraterone in both a time-staggered and non-staggered manner. Abiraterone was administered 24 hours before (top), simultaneously (middle), or 24 hours after (bottom) a second drug. Seventy-two hours after the final drug addition viability relative to control was measured. **(B)** Assessing synergy between the abiraterone and a Plk1 inhibitor (BI2536 4 nM) or microtubule stabilizer (docetaxel 5 nM). C4-2 CRPC cells growing in media containing csFBS were subjected to increasing concentrations of abiraterone in the absence (black line) or presence (red line) of the second drug for 72 hours. The concentration of the second drug was chosen on based on a 20-30% decrease in viability relative to control when used in isolation. Mean ± SEM (n = 3) is shown. Expected viability (dashed black line) was calculated according to the Bliss Independence model of drug additivity. Synergy, depicted by the green area, is a decrease in viability beyond the expected additive effect. **(C-E)** C4-2 cells grown in media containing csFBS were subjected dose matrices of increasing abiraterone (1, 5, 10, and 20 μM) and either BI2536 (C), docetaxel (D), or doxorubicin (E) at concentrations of 1, 10, and 50 nM for BI2536 and docetaxel; 1, 10, and 50 μM for doxorubicin. Drugs were administered in isolation and in all pairwise combinations in a time staggered and simultaneous manner as depicted in (A). The left column of heatmaps are the observed viability measurements from this screen. Heatmap representations of the expected viability based on the Bliss Independence model of drug additivity are in the middle column. Synergy observed in these matrices is depicted in the right column, where the brighter green features indicated more synergy. The data in (B) and (C) are from independent experiments. **(F)** Graphical representation of volumetric measurement of synergy. Viability measurements from a dose matrix can be plotted relative to control in three dimensions to obtain a dose response surface (middle). From this an expected response surface is calculated (top). The expected minus observed response surface (bottom) is calculated and the total synergy present in the dose matrix is calculated as the integrated volume beneath the expected minus observed surface. **(G)** Synergy volume measurements from dose-response matrices in (C-E). Mean ± SEM (n = 3) is shown. **(H)** C4-2 cells in media containing csFBS were treated with 5 nM BI2536, 10 μM abiraterone, and the combination. Viability relative to control was measured daily, mean ± SEM (n = 3) is shown. The expected viability for each timepoint was calculated using the Bliss Independence model of drug additivity. **(I)** C4-2 cells grown in media containing csFBS were subjected to the BI2536 and abiraterone for 72 hrs, fixed, stained with DNA dye SYTO 60, and analyzed by fluorescent imaging. **(J)** Androgen-dependent LNCaP prostate cancer cells were grown in media containing csFBS and subjected to increasing concentrations of abiraterone in the presence (black lines) or absence (red lines) of BI2536 (10 nM). Viability relative to control at 72 hours was measured. Mean values ± SEM (n = 3) are shown. Expected viability (dotted black line) according to the Bliss Independence model of drug additivity is plotted for comparison.

Following drug co-treatment, the observed cell viability was compared to the results expected from simple drug additivity according to the Bliss independence model (BLISS, 1939) (and see Patterson et al., 2019). These experiments revealed dramatic synergy between the Plk1 inhibitor BI2536 and abiraterone (**Figure 1B**), particularly when the entire dose-response matrix is analyzed (**Figure 1C).** By comparison, the expected and observed cell death responses following docetaxel and abiraterone co-treatment essentially overlap (**Figures 1B and D**), as did the responses to doxorubicin and abiraterone (**Figure 1E**). Synergy observed within each dose matrix was quantified by plotting the observed and expected response surfaces and defining synergy as the integrated volume between the observed and expected surfaces (**Figure 1F**) as described previously (53). Clear synergy was observed between the Plk1 inhibitor BI2536 and abiraterone regardless of drug sequencing (**Figure 1G**). Little to no synergy was observed between abiraterone and either docetaxel or doxorubicin in these matrices (**Figure 1G**). A more detailed time course readily confirmed synergy between Plk1 inhibitors and abiraterone, indicating not only suppression of cell growth but a decrease in cell number over time, consistent with cancer cell killing (**Figure 1H**). This was further validated by direct measurement of cell confluence using the SYTO 60 staining (**Figure 1I**).

The C4-2 cell line is a widely used model for CRPC that was derived by passage of LNCaP androgen-dependent prostate cancer cells through castrated mice (54). In contrast to C4-2 cells, co-treatment resulted in additivity but not synergy in the LNCaP parental cell line (**Figure 1J)**. The observation of synergy in C4-2 cells (**Figure 1B**), but not in LNCaP cells (**Figure 1J**), suggests this phenotype could be related to the mechanism(s) by which C4-2 cells acquired a CRPC phenotype and raised the possibility that Plk1 could play a role in maintaining AR pathway signaling in CRPC cells.

### Plk1 inhibitor-abiraterone synergistic killing of CPRC cells is independent of abiraterone’s effects on AR signaling but specific to inhibition of the mitotic kinase Plk1

To assess the relative importance of AR signaling in this synergistic response, we first tested whether abiraterone and Plk1 inhibitors were equally synergistic in non-charcoal-stripped FBS. Not only did synergy between abiraterone and Plk1 inhibitors in C4-2 cells remain, it appeared to be more robust in the presence of exogenous androgens (**Figure 2A**). Furthermore, the presence or absence of androgens did not alter LNCaP cell’s non-synergistic cell viability response to this drug combination (**Figure 1J and Supp. Figure 1B**).

**Figure 2.**
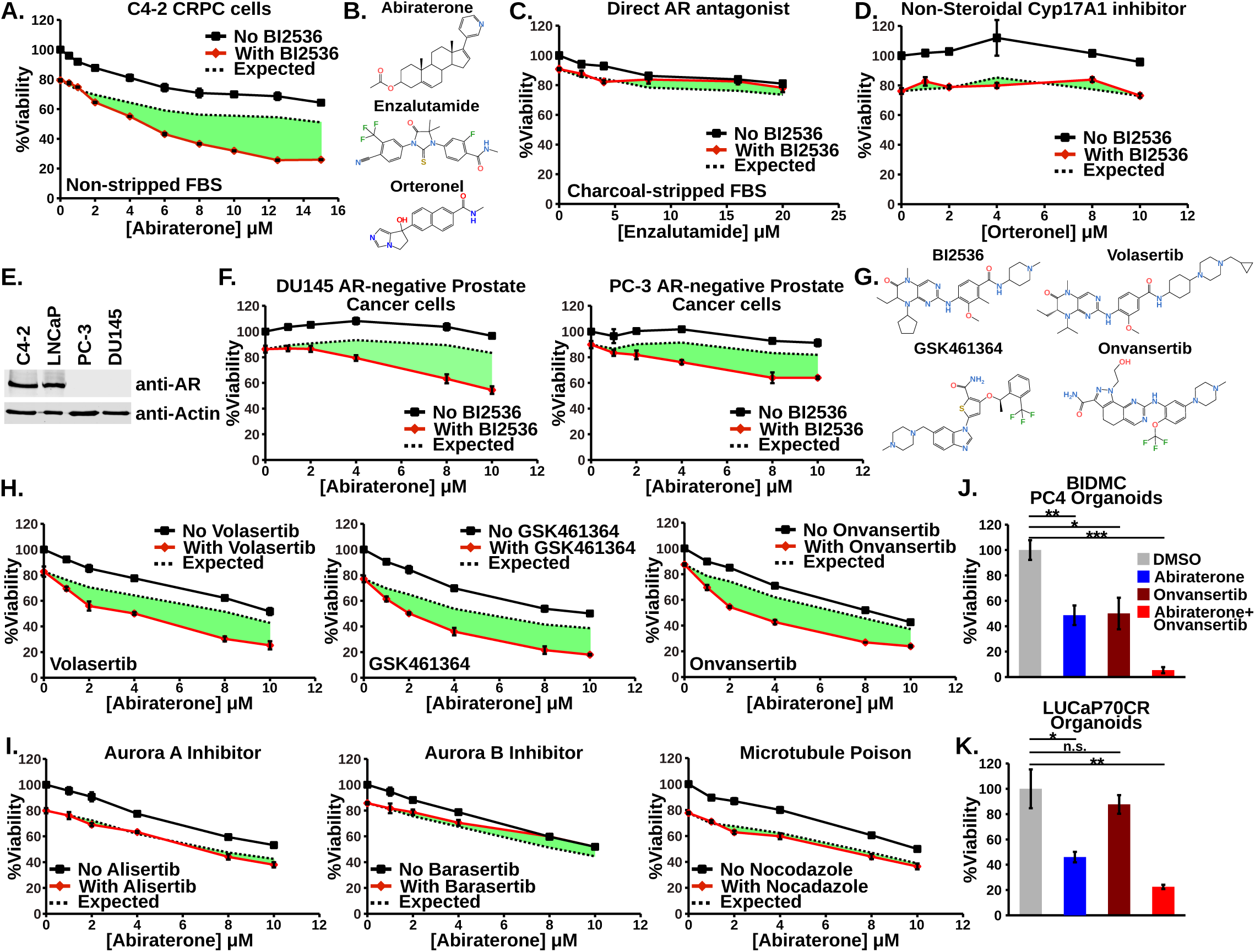
Plk1 inhibitor – abiraterone synergy is independent of AR signaling, occurs in multiple prostate cancer cell lines, and is specific to Plk1 activity. **(A)** C4-2 cells were grown in media containing non-charcoal-stripped FBS and subjected to increasing concentrations of abiraterone in the presence (red line) or absence (black line) of the Plk1 inhibitor BI2536 (3 nM) for 72 hours. Mean ± SEM (n = 3) is shown. Expected viability (dashed black line) was calculated according to the Bliss Independence model of drug additivity. Synergy, depicted by the green area, is a decrease in viability beyond the expected additive effect. **(B)** Comparison of the chemical structures of the antiandrogens abiraterone acetate, enzalutamide, and orteronel (TAK-700). Abiraterone is a pregnenolone derivative and thus shares many chemical features with endogenous steroids. **(C)** C4-2 cells were grown in media containing charcoal-stripped FBS and synergy between the antiandrogen enzalutamide, a direct AR antagonist, and Plk1 inhibitor (BI2536 2.5 nM) was assessed and plotted as in (A). See related figure **Supp. Figure 1C** for similar experiment performed using non-stripped FBS. **(D)** C4-2 cells were grown in media containing FBS and synergy experiments between the nonsteroidal Cyp17A1 inhibitor orteronel (TAK-700) and BI2536 (3 nM) was performed and plotted as in (A). **(E)** Immunoblot of lysates from the indicated cell lines using antibodies directed against full length AR. **(F)** DU145 and PC-3 AR-negative prostate cancer cells plated in media containing FBS and synergy between abiraterone and BI2536 (5 nM and 7.5 nM for DU145 and PC-3, respectively) was measured and analyzed as in (A). **(G)** Chemical structures of the panel of Plk1 inhibitors used. While BI2536 and volasertib share some chemical moieties, both GSK461364 and onvansertib are structurally distinct amongst this panel. **(H)** C4-2 cells were grown in media containing FBS and synergy between abiraterone and the three Plk1 inhibitors volasertib (5 nM), GSK461364 (5 nM), and onvansertib (10 nM) was assessed as in (A). See related figure **Supp. Fig 1D** for similar experiment performed using csFBS. **(I)** C4-2 cells were grown in media containing FBS and subjected to increasing concentrations of abiraterone in the presence or absence of alisertib (20 nM, left), barasertib (200 nM, middle), or nocodazole (200 nM, right). Doses were chosen based on a ∼20% decrease in viability when these antimitotic drugs were used in isolation. Synergy between these drugs was assessed as in (A). **(J, K)** BIDMC PC4 CRPC organoids were treated with onvansertib (200 nM) and abiraterone (10 μM) for 8 days and then relative viability in this 3D culture was assessed. LUCaP70CR CRPC organoids were treated with onvansertib (40 nM) and abiraterone (10 μM) for 5 days and then relative viability assessed n.s. not significant, * p ≤ 0.05, ** p ≤ 0.01, and *** p ≤ 0.001 by Student’s two-tailed t-test

To further explore the extent to which AR signaling inhibition was involved in Plk1 inhibitor-abiraterone synergy, we next tested whether Plk1 inhibitors would synergize with enzalutamide, another clinically used direct AR antagonist that is structurally distinct from abiraterone and does not inhibit Cyp17A1 (**Figure 2B, Supp. Figure 1A**). Surprisingly, no synergy was observed between enzalutamide and Plk1 inhibitors when C4-2 CRPC cells were grown using either csFBS or FBS (**Figure 2C, Supp. Figure 1C**). Under the treatment conditions used, both abiraterone and enzalutamide inhibit AR transcriptional activity to equivalent extents (see below), indicating that the AR inhibition by abiraterone does not drive synergistic killing when this drug is combined with Plk1 inhibitors. Because abiraterone inhibits Cyp17A1, preventing androgen synthesis as well as glucocorticoid and estrogen synthesis, some of the abiraterone AR-independent effects could be due to either loss of other steroid hormones or their accumulation of metabolites upstream of Cyp17A1 (**Supp. Figure 1A**). A non-steroidal Cyp17A1 inhibitor (orteronel/TAK-700) (**Figure 2B**), however, did not synergize with Plk1 inhibitors (**Figure 2D**). These data indicate that neither abiraterone’s effects on AR signaling, or on steroid biosynthesis in general, underlies the synergistic killing observed when the drug is combined with Plk1 inhibition.

In order to conclusively determine that this synergy is independent of AR signaling, we treated both DU145 and PC-3 prostate cancer cell lines with Plk1 inhibitors and abiraterone. PC-3 and DU145 cells lines lack AR expression (**Figure 2E**) and exhibit characteristics associated with a neuroendocrine prostate cancer phenotype (Leiblich et al., 2007; Tai et al., 2011). Despite the lack of AR expression, both DU145 and PC-3 cells demonstrated synergistic killing by the combination of abiraterone and the Plk1 inhibitor BI2536 comparable to that observed in C4-2 cells (**Figure 2F**). Taken together, these data firmly establish that the synergy between abiraterone and Plk1 inhibitors cannot be solely ascribed to suppression of AR signaling, a conclusion that is further reinforced by multiple additional experiments described below.

We next explored whether this abiraterone synergy was truly dependent on Plk1 by examining a panel of Plk1 inhibitors with disparate chemical structures (**Figure 2G**). While BI2536 and volasertib, can inhibit Plk2 and Plk3, GSK461364, and particularly onvansertib are Plk1-specific inhibitors (Gutteridge et al., 2016). All three of these additional Plk1 inhibitors similarly sensitized C4-2 CRPC cells for synergistic killing by abiraterone regardless of the presence or absence of androgens (**Figure 2H, Supp. Figure 1D**).

Plk1 is intimately involved in choreographing multiple stages of mitotic progression, and arrest within mitosis is a well-documented effect of Plk1 inhibition. It was therefore plausible that any drug capable of arresting cancer cells in mitosis would synergize with abiraterone for cancer cell killing. However, as shown previously in **Figure 1B**, co-treatment with the microtubule-stabilizing taxane docetaxel, which causes prometaphase and metaphase arrest, did not sensitize C4-2 cells to abiraterone. To further explore this, we also tested inhibitors of the mitotic kinases Aurora A (alisertib) and Aurora B (barasertib), and inhibition of microtubule polymerization with nocodazole, in combination with abiraterone. As shown in **Figure 2I**, none of these agents caused synergistic tumor cell killing when combined with abiraterone. These data strongly argue that the synergistic anti-tumor effects observed between abiraterone and Plk1 inhibitors involve some unique aspect of Plk1 function rather than a generic effect on mitotic arrest.

Based on these observations, we evaluated the effect of the Plk1 inhibitor onvansertib in combination with abiraterone in two 3-D prostate cancer organoids representative of CRPC. Both the BIDMC PC4 and LuCaP70CR organoid models were generated from CRPC patient-derived xenografts (PDX) (56), express high levels of AR, and grow in androgen-deprived media. As shown in **Figures 2J and K** onvansertib or abiraterone monotherapy inhibits cell growth compared to control, but the combination treatment elicits a significant synergistic response. These results indicate that the phenotype described above is not confined to 2-D models and can be reproduced in 3-D prostate cancer organoids.

### Abiraterone treatment upregulates mitosis and mitotic spindle related gene sets in a synergy-specific and AR-independent manner

To gain insight into the underlying mechanism of abiraterone-Plk1 inhibitor synergistic prostate cancer cell killing, we utilized a comprehensive RNA sequencing experiment to identify transcriptional changes that occur upon abiraterone treatment that are both AR-independent and specific to the Plk1 inhibitor synergistic phenotype (**Figure 3A**). The Plk1 inhibitor onvansertib was used in these experiments because it is highly specific to Plk1 and is actively being investigated in human cancer patients. Enzalutamide was included to separate the AR-dependent and AR-independent transcriptional effects of abiraterone. LNCaP cells were included to identify differential gene expression that was specific to a synergistic phenotype, since LNCaP cells are very similar to C4-2 cells, but do not show synergy (**Figure 1J**). Cells were treated with the indicated drugs for 16 hours in non-stripped serum, then harvested for RNA sequencing. After gene-level quantification, gene set variation analysis (GSVA) was used to transform the data to pathway-level quantification so that the relative expression of biologically meaningful gene sets could be readily compared between the various conditions (57). Differentially expressed gene sets were then identified using limma (58). The relative effects of abiraterone and enzalutamide on AR-dependent transcription was determined using the Hallmark Androgen Response gene set. This curated list of androgen responsive genes has been demonstrated to inform relative AR transcriptional activity (59). The observation that both antiandrogens suppress AR-dependent transcription to an equivalent extent in both cell lines (**Figure 3B**), yet only abiraterone synergizes with onvansertib, and only in C4-2 cells, further supports the notion that this drug combination synergy is not due to suppressed AR signaling. Furthermore, this indicates the utility of using enzalutamide for elucidating the AR-independent effects of abiraterone. Importantly, the small effect onvansertib treatment had on the expression of AR-dependent genes (**Figure 3B**) and AR transcript itself (**Figure S2A**) occurred in both C4-2 and LNCaP cells. Abiraterone treatment did not decrease Plk1 transcript levels (**Figure S2B**). This indicates that the synergistic tumor cell killing seen with this drug combination is not due to inter-regulation of these pathways at the level of transcription.

**Figure 3.**
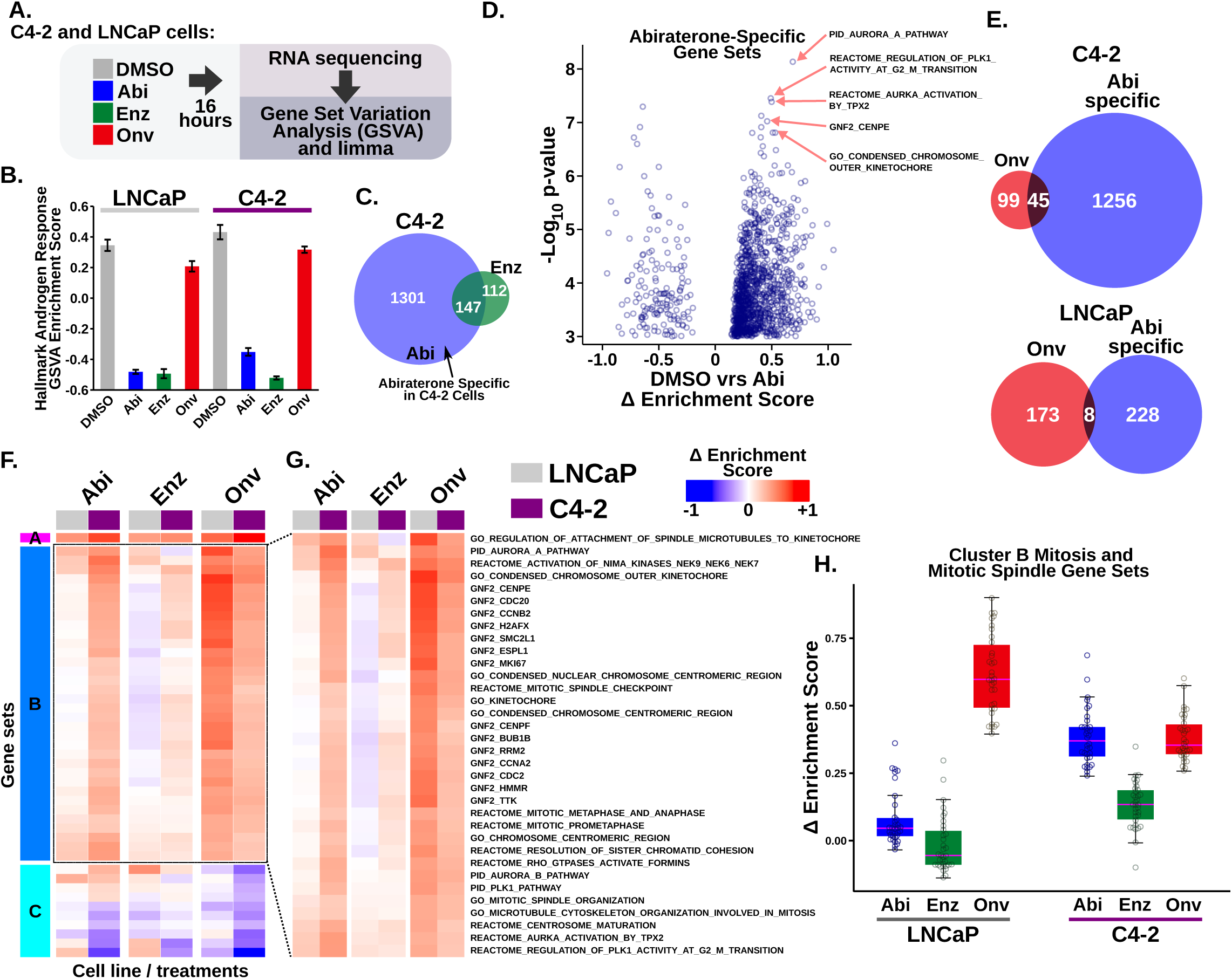
Abiraterone treatment induces an AR-independent synergy-specific gene expression signature dominated by mitosis and mitotic spindle related gene sets. **(A)** Diagram of the RNA sequencing experimental design that was performed in triplicate using LNCaP cells that do not have a synergistic phenotype and C4-2 cells that respond synergistically to the combination of abiraterone and onvansertib. C4-2 and LNCaP cells were grown in media containing FBS and treated with 5 μM abiraterone, 10 μM enzalutamide, and 15 or 30 nM onvansertib, respectively. In this figure abiraterone, enzalutamide, and the Plk1 inhibitor onvansertib are abbreviated Abi, Enz, and Onv, respectively. After alignment of reads and gene-level quantification, GSVA was performed to transform the data matrix into gene set scores. Differentially expressed gene sets were identified using limma by comparing vehicle control to abiraterone, onvansertib, or enzalutamide for both C4-2 and LNCaP cells separately. **(B)** AR-dependent transcription was investigated by focusing on the Hallmark Androgen Response gene set in mSigDB after GSVA analysis. Bars indicate mean ± SEM. Abiraterone and enzalutamide suppress AR-dependent transcription to a comparable extent in this setting. Plk1 inhibitor treatment of cells does not suppress AR-dependent transcription. **(C)** Comparison of differentially expressed gene sets in C4-2 cells treated with abiraterone or enzalutamide. Shown in the Venn diagram is the overlap of significant gene sets (FDR ≤ 0.01) between the two contrasts. There are a substantial number of abiraterone-specific gene sets which are not differentially expressed during enzalutamide treatment. **(D)** Volcano plot of abiraterone-specific differentially expressed gene sets in C4-2 cells. The x-axis is the Δ enrichment score representing the difference in mean GSVA gene set enrichment scores between abiraterone and DMSO treatment, where positive numbers indicate upregulation in abiraterone-treated samples relative to DMSO controls. The most highly significant upregulated gene sets, as indicated by red arrows, are related to mitosis and the mitotic spindle. GNF2 gene sets represent cancer gene neighborhoods (89). The members of these gene sets have expression patterns highly covariant with the seed gene listed in the name. **(E)** Comparison of abiraterone-specific and onvansertib differential gene set expression in C4-2 and LNCaP cells. Shown is the overlap of significant gene sets (FDR ≤ 0.01) between these contrasts in the two cell lines. Onvansertib and abiraterone share a large number of significant gene sets only in C4-2 cells. **(F)** Unsupervised hierarchical clustering was performed using all treatments (columns) and the 45 gene sets (rows) that were common amongst gene sets identified in (E) in C4-2 cells. Columns depicting LNCaP and C4-2 gene set expression are marked at the top with grey and purple boxes, respectively. For expanded heatmap that includes all row labels see **Supp. Figure 2C**. From this we identified three clusters labeled A-C. **(G)** Expanded view of cluster B gene sets that are induced by abiraterone but not enzalutamide in C4-2 but not LNCaP cells, organized and displayed as in (F). Nearly all of the gene sets in cluster B are related to mitosis or the mitotic spindle. **(H)** Box and whisker plot for direct comparison of abiraterone, enzalutamide, and onvansertib’s effects on gene sets in cluster B in both C4-2 and LNCaP cells. Y-axis indicates the differences in gene set expression between vehicle control and the indicated drug treatment (Δ enrichment score). Individual gene sets are overlaid as circles.

To explore transcriptional effects of abiraterone that go beyond inhibition of AR signaling, we sought to identify an abiraterone-specific expression signature defined as gene sets that were differentially regulated by abiraterone, but not enzalutamide in C4-2 cells. Interestingly, a large number of gene sets were found to be uniquely differentially expressed after abiraterone treatment (**Figure 3C**). Conversely, the majority of the gene sets that were differentially regulated upon enzalutamide treatment were gene sets that were also altered by abiraterone treatment. When we examined the composition of the abiraterone-specific gene sets it was notable that, among the most statistically significant gene sets that were upregulated following abiraterone treatment, several were related to mitosis and Plk1 function, though without affecting Plk1 mRNA levels themselves (**Figure 3D**). This observation suggests that the abiraterone-specific effects responsible for synergistic killing of C4-2 cells may functionally overlap with the consequences of Plk1 inhibition. We, therefore, next identified gene sets that were both abiraterone-specific and differentially regulated by onvansertib treatment in C4-2 and LNCaP cells (**Figure 3E**). Among gene sets that were differentially expressed after onvansertib treatment in C4-2 cells, roughly a third were also differentially expressed after abiraterone exposure. This was in sharp contrast to LNCaP dataset that contained very few gene sets shared between those differentially regulated after abiraterone or onvansertib treatment. We focused on the 45 differentially expressed gene sets that were both abiraterone-specific and significantly altered by onvansertib treatment in the C4-2 cells that showed a synergistic response (**Figure 3E**) and further analyzed them by unsupervised hierarchical clustering. This resulted in three clusters (Clusters A-C, **Figure 3F and Supp. Figure 2C**). Cluster B, the largest of the three clusters, was composed entirely of gene sets related to mitosis or mitotic spindle assembly (**Figure 3G**). The gene sets within Cluster B were then mapped back onto both the LNCaP and C4-2 drug treatment/RNA expression data revealing that these gene sets were specifically upregulated by both abiraterone and onvansertib treatment in C4-2 cells, which showed synergistic killing, but only by onvansertib in the LNCaP cell line, which did not demonstrate synergistic cell death (**Figure 3H**). Importantly, these gene sets were not significantly upregulated by enzalutamide in either cell line. Taken together, this abiraterone-specific and synergy-specific gene set signature indicates that abiraterone has AR-independent effects that result in upregulation of genes related to mitotic progression, and further suggests that abiraterone may have direct effects on the process of mitosis itself.

### Abiraterone treatment causes multiple mitotic defects in a manner that is distinct from its AR blockade effects

The observation that abiraterone treatment resulted in an AR-independent induction of mitosis- and spindle-related gene sets suggests that abiraterone may have direct effects on mitotic progression or mitotic spindle assembly. To investigate this, we examined the morphology of mitotic spindles after abiraterone treatment by indirect immunofluorescence. C4-2 CRPC cells were treated with DMSO abiraterone, and enzalutamide for 24 hours, fixed and then stained with antibodies against tubulin and the centromeric histone CENP-A, as well as DAPI to visualize the mitotic chromosomes. In contrast to treatment of C4-2 cells with Plk1 inhibitors or other mitotic spindle poisons (53), abiraterone did not dramatically alter spindle organization when used at doses that elicit synergy. Close examination of mitotic spindles, however, revealed an altered chromatin morphology in which abiraterone treatment resulted in less compact chromosomes during metaphase. In both DMSO control and enzalutamide-treated cells, chromosomes aligned at the metaphase plate contain highly condensed chromatin such that discrete chromosome arms are readily observable. Abiraterone treatment, however, resulted in the majority of metaphase cells having insufficiently condensed chromatin to define individual chromosome arms (**Figure 4A-B and Supp. Figure 3A**). This is most clearly seen in **Supp. Movies 1, 2, and 3** using deconvolution microscopy to compare the Z-stacked images of DMSO, abiraterone, and enzalutamide treated cells. Decreased chromatin condensation was also accompanied by occasional misaligned chromosomes in abiraterone treated cells. To quantify this defect in chromatin condensation we used CENP-A staining to directly measure the intercentromeric distance among mitotic sister chromatids (**Supp. Movie 4**), a well-accepted method for documenting defects in chromatin condensation during mitosis (60, 61). This revealed that abiraterone treatment, but not enzalutamide treatment, increased the intercentromeric distance between sister chromatids (**Figure 4C and D**), consistent with a defect in chromosome condensation in abiraterone-treated cells.

**Figure 4.**
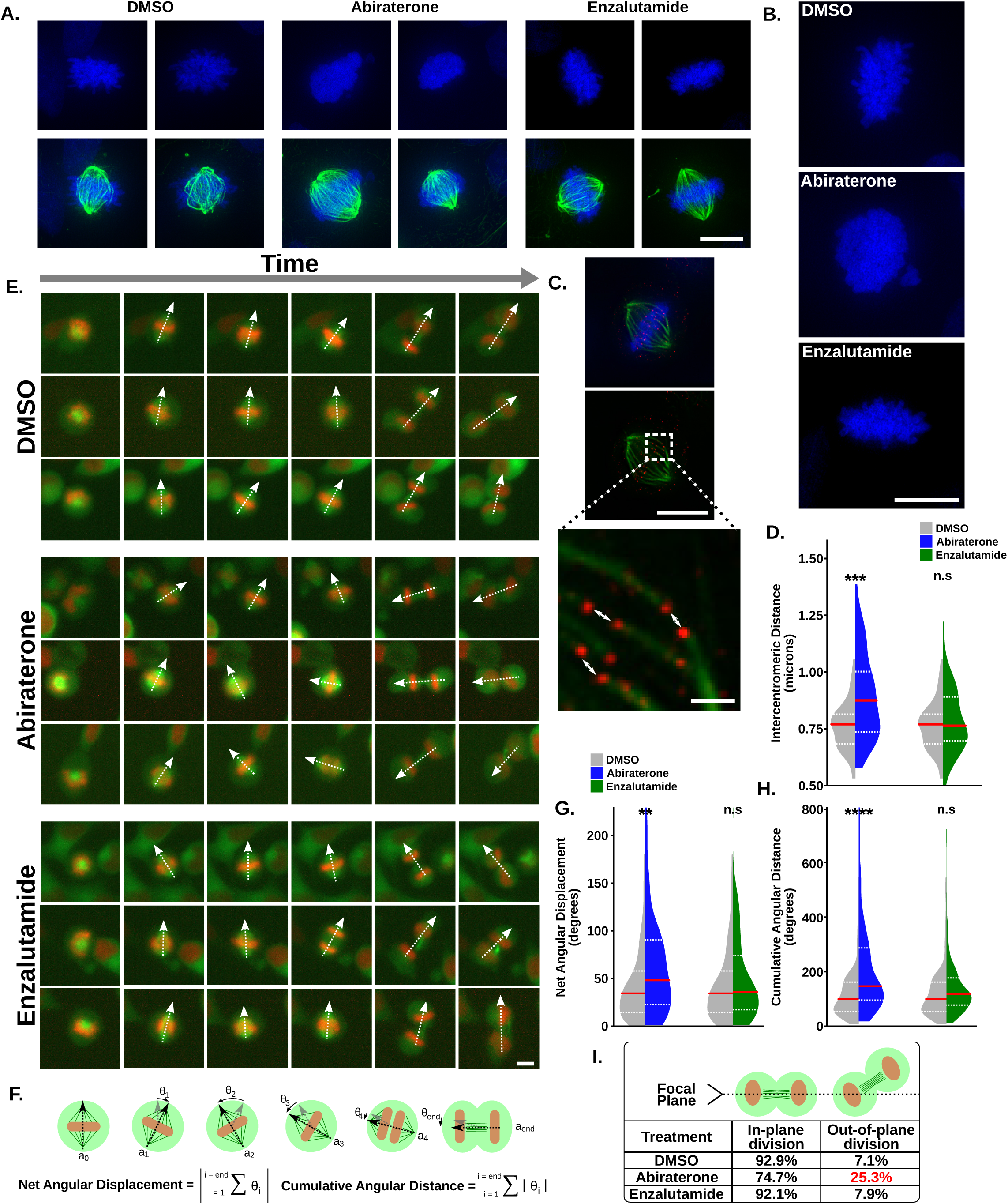
Abiraterone treatment prevents stable mitotic spindle orientation and impairs chromosome condensation independently of effects on AR signaling. **(A)** C4-2 CRPC cells were treated with DMSO, abiraterone (5 μM), or enzalutamide (10 μM) for 24 hours prior to fixation and staining with DAPI (blue) for visualization of mitotic chromosomes as well as antibodies against tubulin (green) to examine mitotic spindle structure. Shown are two examples each with additional examples in **Supp. Figure 3A**. Note the well-defined chromosome arms in the DMSO and enzalutamide treated cells in contrast to the ill-defined and apparently decondensed chromosome arms seen in abiraterone treated cells. Scale bar lower right represents 10 μm. See **Supp. Movies 1, 2 and 3** for comparison of deconvolved Z-stacked images. **(B)** Enlarged micrographs of cells treated and analyzed as in (A) showing decondensed chromatin. Scale bar lower right represents 10 μm. **(C)** C4-2 cells treated as in (A) were stained for the centromeric histone CENP-A (red). Centromeres on sister chromatids were identified on the basis of their paired orientation in three-dimensional space as well as microtubule fibers emanating from opposite poles of the spindle. See **Supp. Movie 4** for more information on identification of centromeres on sister chromatids. Intercentromeric distance was measured as the distance between paired centrosomes (white arrows). Scale bar lower right of top images represents 10 μm. Scale bar lower right of bottom represents 1 μm. **(D)** Violin plots depicting the distribution of intercentromeric distances observed in DMSO, abiraterone and enzalutamide treated cells. Intercentromeric distance was measured for five pairs of centromeres identified in ten cells per condition (n=50). Red line represents the median and dotted white lines are upper and lower quartiles. *** p ≤ 0.001, n.s. not significant using a two-tailed Mann-Whitney U test. **(E)** C4-2 cells expressing H2B-mCherry and mEmerald-tubulin were treated with vehicle control, abiraterone (5 μM), or enzalutamide (10 μM), and analyzed by time-lapse live-cell microscopy. Individual cells (rows) were tracked through time as they progress from prophase (frame 1), as judged by a single cell with condensed chromatin, to cytokinesis (frame 6), as judged by the presence of two daughter cells containing decondensed chromatin. Images were acquired at 15-minute intervals. The dashed white arrows represent the major axis of the mitotic spindle (from pole to pole). White bar at bottom right represents 10 μm. Related to **Supp. Movie 5.** **(F)** Mitotic cells were analyzed and the vector axis (a_i_) of the mitotic spindle was tracked over time from when it was first apparent until telophase. Frames are denoted by the subscript i corresponding to the frame number starting at zero. The rotation between each frame was then calculated (grey to black arrows; θ_i_) where clockwise rotation was considered positive and counterclockwise rotation negative. Net angular displacement was defined as the absolute value of the sum of all stepwise rotations from start to finish, and indicates the final amount of rotation that occurred between the initial and final angle. Cumulative angular distance was calculated as the sum of the absolute values of all stepwise rotation angles, and indicates the total amount of spindle rotation that occurred throughout mitosis. **(G, H)** Violin plots comparing the distribution of net angular displacement and cumulative angular distance in DMSO versus abiraterone or enzalutamide treated cells. Red line represents the median and dotted white lines are upper and lower quartiles (n ≥ 92). ** p ≤ 0.01, **** p ≤ 0.0001, n.s. not significant using a two-tailed Mann-Whitney U test. **(I)** During the mitotic spindle angle analysis described in (E) the frequency of mitotic divisions that occurred either parallel or not parallel to the growth surface in the three different conditions were also tracked. Nonparallel division was apparent when one daughter cell left the focal plane during cytokinesis. See cartoon above table as well as **Supp. Movies 6 and 7**.

We next sought to examine the impacts of abiraterone treatment on the dynamic process of mitosis by tracking living cells over time. C4-2 cells expressing histone H2B-mCherry and mEmerald-tubulin were analyzed by time-lapse microscopy and both overall mitotic progression and spindle formation were assessed during abiraterone or enzalutamide treatment. Abiraterone, but not enzalutamide, caused a statistically significant but small increase in the duration of mitosis (**Figure Supp. 4B**). This finding prompted us to perform a more detailed analysis of mitotic spindle dynamics, which further revealed significant defects in mitotic spindle orientation caused by abiraterone, but not by enzalutamide. Spindles in abiraterone-treated cells failed to form a stable axis of cell division with respect to the cell cortex, and instead displayed continual rotation as the cells progressed from prometaphase through telophase (**Figure 4E, Supp. Movie 5**). The angle of the spindle’s major axis was tracked relative to its initial position, from the time when it was first apparent until telophase. This allowed calculation of both the net difference and cumulative amount of rotation that occurred throughout the entire mitotic process (termed displacement and distance, respectively; **Figure 4F**). A clear increase in the net displacement and total rotational angle was observed during mitosis in abiraterone-treated cells (**Figures 4G-H**). No such changes in spindle orientation during mitosis were observed in enzalutamide-treated cells, indicating that it is not a consequence of AR-inhibition. Notably, the increase in spindle rotation occurred throughout all stages of mitosis, and was not a consequence of prolonged mitosis, spindle collapse or deterioration (**Supp. Figure 3C**). Consistent with impaired spindle orientation abiraterone treatment caused an increase in the frequency of cells undergoing out-of-plane cell division, i.e. non-parallel to the plane of growth (**Figure 4I, Supp. Movie 6 and 7**).

### Abiraterone in combination with Plk1 inhibition causes synergistic mitotic arrest followed by cell death in a spindle assembly checkpoint-dependent manner

The findings described above, as portended by our RNA sequencing analysis, indicates that abiraterone exerts multiple AR-independent effects on the process of mitosis that, while falling short of complete inhibition of cell division, could render a subset of cancer cells more susceptible to inhibition of Plk1. To examine this hypothesis, C4-2 CRPC cells were treated with abiraterone, onvansertib, or the combination, and analyzed by flow cytometry for their cell cycle distribution by staining with the DNA dye DAPI, and with antibodies against phospho-serine 10 histone H3 (pHH3), a mitotic marker. Abiraterone caused a small but statistically significant increase in the percentage of mitotic cells, while onvansertib markedly increased pHH3+ population consistent with mitotic arrest (**Figure 5A, B**). Interestingly, the combination of onvansertib plus abiraterone induced a two-fold increase in mitotic arrest in comparison to onvansertib single agent, suggesting that abiraterone’s effects on mitosis likely contribute to the enhanced accumulation of mitotic cells following Plk1 inhibitor co-treatment.

**Figure 5.**
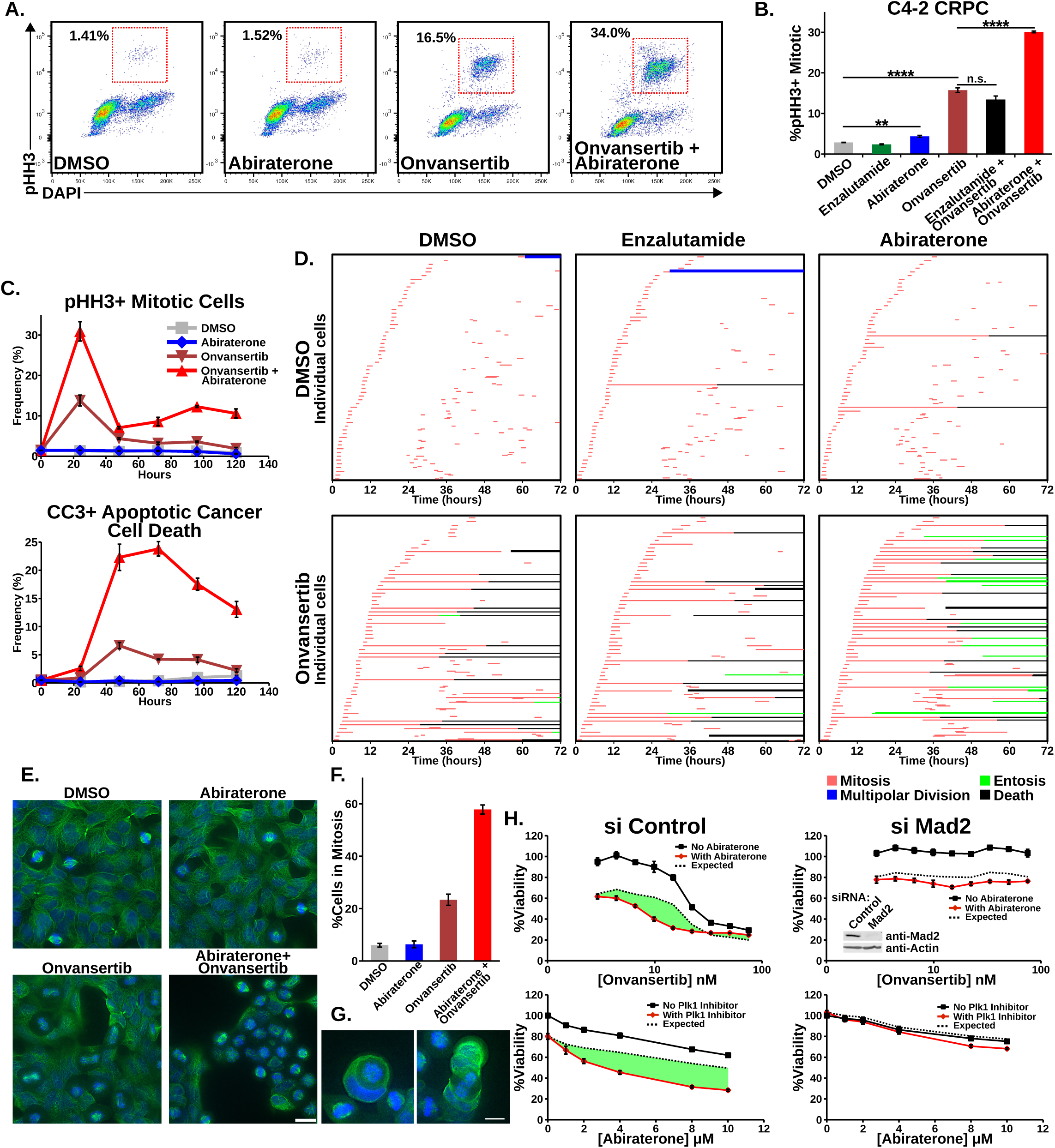
The combination of abiraterone and Plk1 inhibitors cause synergistic mitotic arrest followed by tumor cell death in a SAC-dependent manner. **(A)** C4-2 CRPC cells were subjected to DMSO, abiraterone (5 μM), onvansertib (15 nM), or the combination for 24 hours, fixed, and cell cycle distribution determined by staining with DAPI and antibodies against phospho Ser10 histone H3 (pHH3) a marker for mitosis. Red boxes indicate the location of mitotic cells in these scatter plots. **(B)** Mitotic arrest was measured as in (A) by flow cytometry in fixed C4-2 cells after 16-hour treatment with abiraterone (5 μM), enzalutamide (10 μM), and onvansertib (15 nM) and the indicated combinations. Mean ± SEM (n = 3) values are shown. n.s. not significant, * p ≤ 0.05, ** p ≤ 0.01, and **** p ≤ 0.0001 by Student’s two-tailed t-test; red arrows indicated the direction of the change. **(C)** C4-2 cells were treated and collected as in (A) over a 120-hour time course. The percentage of mitotic cells (top) was measured by flow cytometry. The percentage of cells in other stages of the cell cycle in this analysis is presented in **Supp. Figure 4C**. Additional staining with antibodies against CC3 was used to determine the percentage of cells undergoing apoptosis (bottom). Mean ± SEM (n = 3). **(D)** Time-lapse microscopy of C4-2 cells expressing H2B-mCherry and mEmerald-tubulin treated with vehicle control, abiraterone (5 μM), and enzalutamide (10 μM); alone and in combination with onvansertib (15 nM). For each condition, 60 cells were analyzed to determine both the duration of mitosis and the cellular phenotypes associated with mitotic arrest. Each row begins by tracking an individual cell until initiation of mitosis (red), the two daughter cells were then tracked which themselves could go through mitosis (red) to create up to four daughter cells. Prolonged mitotic arrest was often followed by cell death (black) or entosis (green). See **Supp. Figure 4D** for additional information on interphase lengths as well as **Supp. Movies 6, 8 and 9**. **(E)** AR-negative DU145 prostate cancer cells were treated with the onvansertib (10 nM) and abiraterone (10 μM) for 12 hours, fixed, and examined by microscopy to determine the frequency and fate of mitotic cells. Tubulin localization was assessed by immunofluorescence (green) and DNA content using DAPI (blue). Scale bar at bottom right indicates 20 μm. **(F)** Quantification of the frequency of mitotic DU145 cells from the microscopy analysis presented in (E). Mean ± SEM values are shown. **(G)** Entosis of DU145 mitotic cells after cotreatment with abiraterone and onvansertib. Scale bar at bottom right indicates 10 μm. **(H)** Disabling the SAC by Mad2 siRNA mediated knockdown eliminated synergy between abiraterone and onvansertib. C4-2 cells were transfected with a control siRNA (N1, left panels) or an siRNA targeting Mad2 (right panels). After 48 hours the cells were replated, and the following day assessed for synergy both in terms of sensitivity to onvansertib in the presence of 8 μM abiraterone (top) and sensitivity to abiraterone in the presence of 15 nM onvansertib (bottom). Shown is viability relative to control 72 hours after drug addition, mean ± SEM (n = 3). The dotted black line represents the expected response based on the Bliss independence model of drug additivity. The immunoblot inset in the upper right panel confirmed Mad2 knockdown 48 hours after siRNA transfection.

To determine whether this synergistic mitotic arrest was truly an AR-independent effect of abiraterone, C4-2 cells were treated with vehicle, abiraterone, or enzalutamide; alone or in combination with onvansertib, and analyzed as above. As shown in **Figure 5B**, the combination of enzalutamide and onvansertib showed essentially the same extent of mitotic arrest as that obtained with onvansertib treatment alone, whereas the abiraterone plus onvansertib combination resulted in synergistic mitotic arrest. Furthermore, siRNA knockdown of Cyp17A1 also failed to enhance mitotic arrest caused by Plk1 inhibition, indicating synergistic mitotic arrest was not due to inhibition of androgen synthesis or accumulation of steroidal hormones upstream of Cyp17A1 (**Supp. Figure 4A and B**). When assessed as a function of time, mitotic arrest caused by this combination peaked at 24 hours and was followed by synergistic induction of apoptotic cancer cell death as judged by cleaved caspase-3 (CC3) staining (**Figure 5C and Supp. Figure 4C**). These data indicate that the AR-independent effects of abiraterone synergize with Plk1 inhibition to cause enhanced mitotic arrest followed by markedly increased apoptosis consistent with mitotic catastrophe (62).

To further validate this conclusion, acquire a more granular understanding of this synergistic mitotic phenotype, and assess cell fate following mitotic arrest, time-lapse microscopy of C4-2 cells expressing H2B-mCherry and mEmerald-tubulin was performed. During a 72-hour time course cells were treated with vehicle, abiraterone, or enzalutamide; both alone and in combination with onvansertib. Individual cells were tracked to analyze both the time spent in mitosis and the consequences of drug treatment on cell division (**Figures 5D**). At the dose of onvansertib used in these experiments, Plk1 inhibition elicited a heterogeneous response, with a subset of cells displaying prolonged mitotic arrest (∼24 hours) while other cells progressed through mitosis at a rate similar to those treated with vehicle alone (**Figure 5D, left top and bottom panels**). Prolonged mitotic arrest was frequently followed by cell death (**Figure 5D and Supp. Movie 8**). The combination of abiraterone and onvansertib markedly increased the proportion of cells that suffered severe mitotic arrest compared to onvansertib treatment alone (**Figure 5D, compare bottom left and right panels**), whereas enzalutamide treatment had no effect on the extent of Plk1 inhibitor induced mitotic arrest (**Figure 5D, bottom left and middle panels)**. While the prolonged mitotic arrest caused by the combination of abiraterone and onvansertib was frequently followed by cell death, we additionally observed a marked increase in entotic cancer cell death (**Figure 5D and Supp. Movie 9**). Entosis is an alternate form of cell death that involves one cell invading another, and has been reported to be a consequence of mitotic arrest in certain situations (62, 63). While entosis occurred among C4-2 cells treated with onvansertib alone at a low frequency, it appeared to play a major role in the manner of cell death among cells treated with the combination of abiraterone and onvansertib. None of these phenotypes were recapitulated by the combination of enzalutamide with onvansertib. To further confirm that the combination of abiraterone and Plk1 inhibitors synergistically disrupts mitosis in prostate cancer cell lines in an AR-independent manner, the AR-negative prostate cancer cell line DU145 that had a synergistic cell death phenotype (**Figure 2F**) was treated with onvansertib, abiraterone, and the combination, and then analyzed by both flow cytometry and microscopy. Despite completely lacking detectable AR protein (**Figure 2E**), these cells demonstrated synergistic mitotic arrest upon treatment with the abiraterone plus onvansertib combination when quantified by flow cytometry (**Supp. Figure 4E**) or through quantification of mitotic figures by microscopy (**Figures 5E and F**). Moreover, entotic figures were also observed among cells treated with both drugs indicating this phenomenon occurs independently of abiraterone’s effects on AR signaling **(Figures 5G).**

Our observation that abiraterone impacts multiple processes in mitosis rendering some cancer cells more susceptible to prolonged mitotic arrest following Plk1 inhibition implies that the synergy we observe between these drugs should depend on a functional spindle assembly checkpoint (SAC). The SAC inhibits progression to anaphase through inhibition of APC/C-Cdc20 until the spindle has achieved chromosome biorientation though proper microtubule-kinetochore attachments and tension between opposite spindle poles (64). Mad2 is an essential component of the SAC, and loss of this protein prevents mitotic arrest caused by spindle assembly defects. When we examined Plk1 inhibitor-abiraterone synergy in C4-2 cells in which Mad2 had been knocked down by siRNA, the synergy was entirely eliminated (**Figure 5H**). Taken together, these data indicate that abiraterone has AR-independent effects that result in multiple defects in mitosis and cell division, creating a vulnerability that can then be targeted by inhibiting Plk1, resulting in synergistic mitotic arrest followed by cancer cell death in a SAC-dependent manner.

### The abiraterone/Plk1 inhibitor combination synergistically kill cancer cells from diverse tumor types in culture and *in vivo*

Given the observation that abiraterone sensitizes a subset of prostate cancer cells to Plk1 inhibition through synergistic disruption of mitosis and subsequent cancer cell death in an entirely AR-independent manner, we considered the possibility that the potency of this combination was not limited to prostate cancer cells. We therefore examined a set of acute myeloid leukemia (AML), pancreatic cancer, and ovarian cancer cell lines for their response to abiraterone, onvansertib, and the combination. As show **Figures 6A and B**, MV-4-11 AML cells, PSN-1 pancreatic cancer cells, and SK-OV-3 ovarian cancer cells showed clear synergistic killing by the combination, while OCI-AML-3 AML, Panc 10.05 pancreatic cancer cells, and OAW28 ovarian cancer cells did not. Notably, it was also generally the case that cell lines that showed synergistic killing also displayed synergistic mitotic arrest as a result of combination treatment, as reflected by the large fold increase in mitotic cells following treatment with abiraterone and onvansertib compared to onvansertib alone (purple bars in **Figure 6C).** None of the non-prostate cancer cell lines expressed detectable levels of the androgen receptor (**Figure 6D**), clearly demonstrating that synergistic mitotic arrest and cell death caused by the combination of abiraterone and Plk1 inhibition is independent of AR signaling.

**Figure 6.**
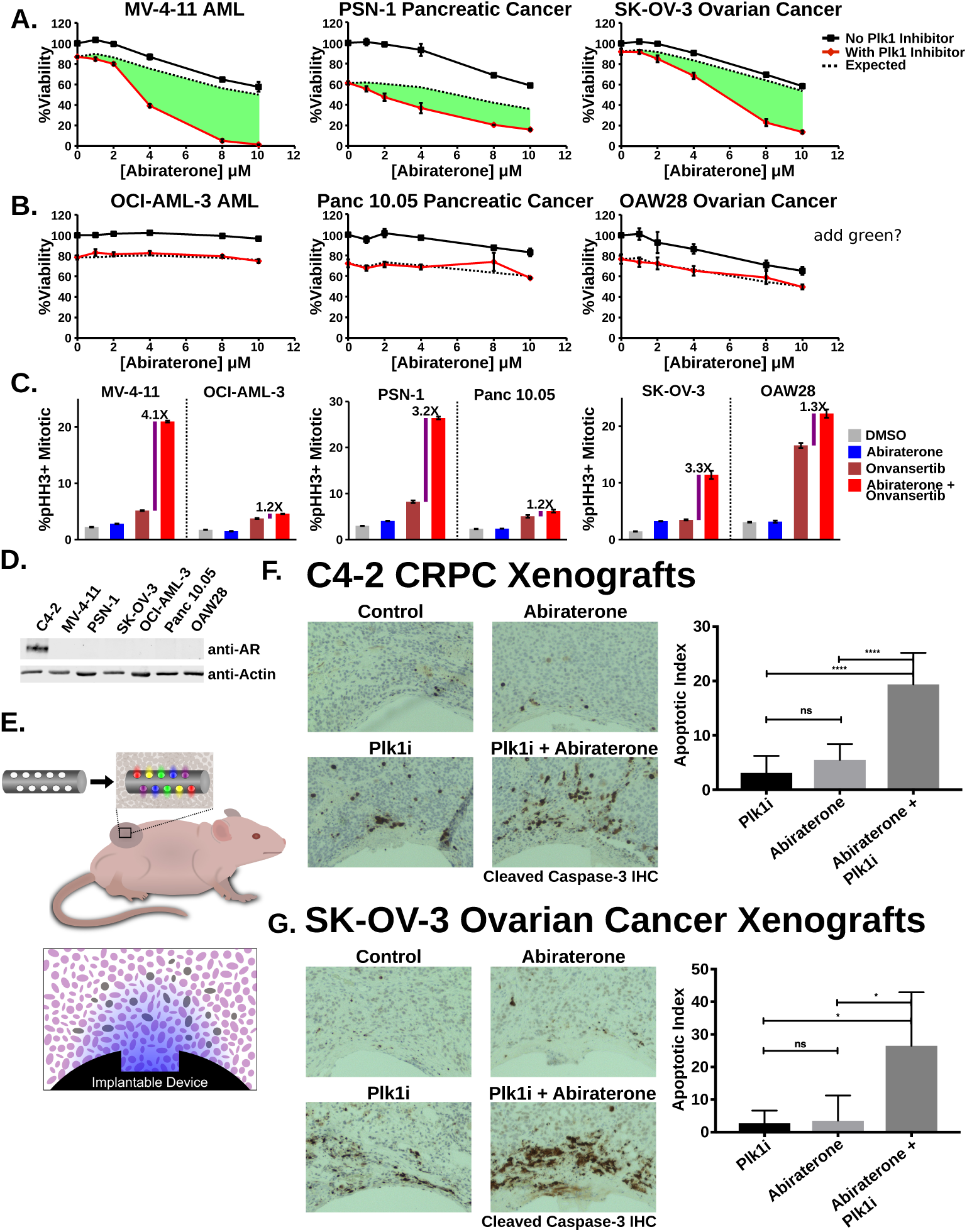
Synergistic mitotic arrest and cancer cell killing by combined abiraterone and Plk1 inhibitor treatment occurs in multiple AR-negative types of cancer cell lines *in vitro* and *in vivo*. (A, B) Six non-prostate cancer cell lines from three indications were tested for synergy between abiraterone and onvansertib. Three of these cell lines have synergistic responses (A) to the combination of abiraterone and onvansertib whereas three do not (B). Each cell line was subjected to increasing concentrations of abiraterone in the absence (black lines) or presence (red lines) of onvansertib (MV-4-11 22.2 nM, OCI-AML-3 22.2 nM, PSN-1 22.2 nM, Panc 10.05 33.3 nM, SK-OV-3 33.3 nM, and OAW28 15 nM). Viability relative to control was measured after 72 hours (mean ± SEM, n = 3). The dotted line represents expected combined result according to the Bliss independence model of drug additivity and the areas shaded in green indicate synergy. **(C)** Cells used in A and B were tested for synergistic mitotic arrest during abiraterone-onvansertib cotreatment by pHH3 staining and flow cytometry. Shown are bar graphs, paired by indication, displaying the percentage of cells that were mitotic after treatment with the indicated drugs for 16 hours. Purple bars indicate the fold-change in percentage pHH3+/mitotic cells between onvansertib treatment alone and the combination of abiraterone and onvansertib. Quantification of the percentage of cells in other stages of the cell cycle is presented in **Supp. Figure 5A**. Abiraterone treatment was 10 μM for all cell lines except OAW28, which was 5 μM. Onvansertib concentrations used were 20 nM for MV-4-11, OCI-AML-3, and PSN-1; 30 nM for Panc 10.05 and SK-OV-3; 15 nM for OAW28. Mean ± SEM (n = 3) values are shown. **(D)** Immunoblot for detection of AR protein in both C4-2 cells and the non-prostate cancer cells used for synergy analysis. **(E)** Diagram of the tumor-implantable device used to examine synergy between abiraterone and Plk1 inhibitors *in vivo*. Colored gradients represent drug concentration gradients created upon device implantation. On the left is the implantable device with respect to the tumor bearing mouse, on the right is a depiction of a single well and drug concentration gradient corresponding to the fixed and stained tumor sections shown in (F) and (G). **(F)** C4-2 CRPC tumors were grown in non-castrated male NCR nude mice and implanted with devices. Microwells were loaded with drug at 12.5% by weight in isolation or each at 12.5% by weight when combined. Fixed tumor sections adjacent to the microwells containing the indicated drugs were stained with antibodies directed against CC3 to detect apoptotic cells (representative examples, left). Clear sections at the bottom of each image are where the implantable device was previously located. On the right is quantification of the percentage of apoptotic cells within a 400 μm radius of wells. Bars represent mean ± SEM where n ≥ 9; n.s. not significant and **** p ≤ 0.0001 by a Student’s two-tailed t-test. **(G)** SK-OV-3 ovarian cancer tumors were grown in female NCR nude mice and implanted with devices. Tumors were processed, analyzed, and quantified as in (F) where n ≥ 5; n.s. not significant and * p ≤ 0.05 by Student’s two-tailed t-test.

To directly examine the potentially greater-than-additive effects of these drugs on apoptotic cancer cell death *in vivo*, we examined human C4-2 CRPC xenografts grown in non-castrated mice using a tumor implantable device capable of achieving multiplexed drug sensitivity testing within tumors(53, 65)(**Figure 6E**). The device consists of a small cylinder containing multiple drug-loaded microwells which is implanted into an existing human tumor grown on the hind flank of a mouse. Localized intratumor release of the drugs or drug combinations generates spatially distinct concentration gradients in the adjacent section of the tumor. Following incubation for multiple days, the tumors are then excised, fixed, sectioned, and stained for CC3 as a marker of apoptotic cancer cell death. As shown in **Figure 6F**, low levels of apoptotic cell death were observed in C4-2 tumor sections adjacent to wells containing abiraterone or the Plk1 inhibitor alone. In contrast, significant apoptotic cell death was seen in regions of the tumor adjacent to wells containing the abiraterone/Plk1 inhibitor combination. The apoptotic index was 3.6-fold higher in fields of cells adjacent to the combination relative to those subjected to abiraterone alone (**Figure 6F**). Because localized delivery of abiraterone within the tumor does not eliminate synthesis of androgens in the testes, this indicates that synergistic tumor killing *in vivo* can still occur in the presence of systemic androgens.

We next used this implantable device to examine drug responses in SK-OV-3 ovarian cancer xenografts, since this AR-negative tumor cell line had demonstrated a pronounced synergistic response when tested in culture (**Figures 6A**). Like the CRPC cells, these ovarian cancer cells showed little apoptotic cell death in response to treatment with either drug alone, but 8.3-fold increase in apoptotic cells in response to treatment with the abiraterone/Plk1 inhibitor combination relative to abiraterone alone (**Figure 6G**). This *in vivo* synergy seen in AR-negative non-prostate cancer tumor xenografts further supports the conclusion that abiraterone has effects on cancer cells that are independent from AR signaling yet synergize with Plk1 inhibition to induce cancer cell death.

### Plk1 inhibition in combination with abiraterone acts synergistically in patient-derived CRPC tumor xenografts

To more closely model the pharmacokinetics and tumor biology present in mCRPC patients, we examined the effects of systemic dosing using a PDX model of CRPC, LVCaP-2CR, grown on the hind flanks of castrated male mice. This PDX expresses the AR-v7 splice variant and is thus relatively resistant to antiandrogens including abiraterone (66). As shown in **Figure 7A** compared to the vehicle only control, each of the single agents had modest effects on tumor growth. The combination of abiraterone and onvansertib, however, reduced tumor growth to a much greater extent. These human PDX tumor responses were analyzed using a modified version of Bliss independence adapted to tumor volume data (67), and the abiraterone/Plk1 inhibitor combination was confirmed to be synergistic *in vivo* (**Figure 7B**).

**Figure 7.**
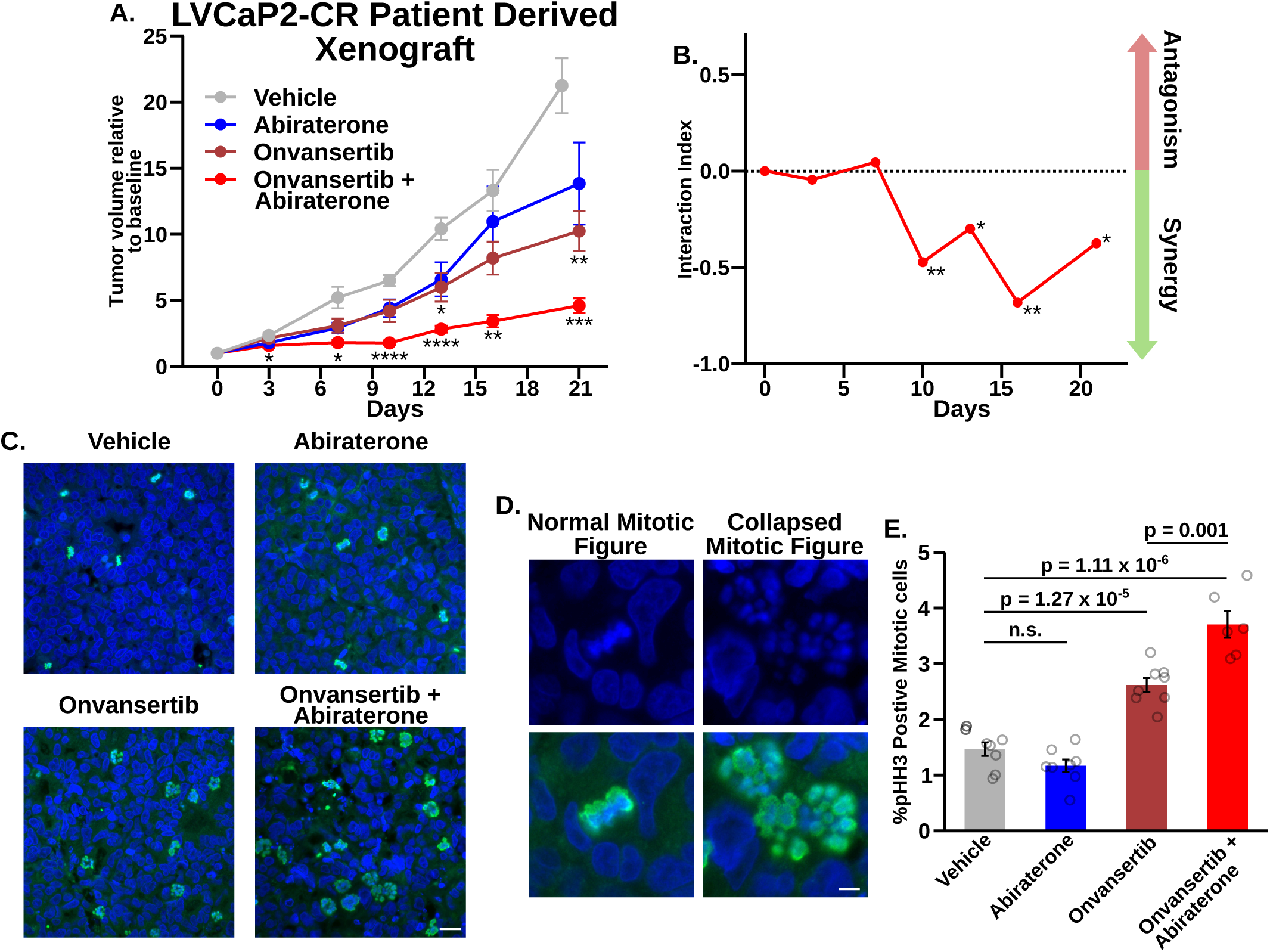
Abiraterone and Plk1 inhibitor synergize *in vivo* and cause enhanced mitotic arrest in a murine PDX model of prostate cancer. **(A)** Castrated male NOD scid gamma mice carrying LVCaP-2CR PDXs were administered the indicated drugs for 21 days. Mice were administered drugs for five consecutive days followed by 24- to 48-hour drug holidays over the course of the experiment. Tumor volume was normalized to the day 0 volume on an individual animal basis. Plotted is the mean ± SEM where n = 6 to 8; *, **, and *** indicate p ≤ 0.05, 0.01, and 0.001, respectively, when comparing the indicated treatment to the vehicle control by Tukey’s multiple comparisons test. **(B)** Synergistic responses were assessed according to (67) where interaction indices below zero indicate a synergistic response. * and ** indicate p ≤ 0.05 and 0.01, respectively, using a bootstrap-t method with 10,000 random samples. **(C)** End-of-study tumors were fixed and sections were stained with DAPI (blue) and antibodies against pHH3 (green). Scale bar at bottom right indicates 20 μm. **(D)** Examples of both normal and abnormal mitotic figures observed in stained tissue sections. Top and bottom micrographs depict DAPI or both DAPI and pHH3 staining, respectively. Abnormal mitotic figures contained mislocalized and decondensed chromatin that resemble collapsed spindles seen after prolonged mitosis by Plk1 inhibition or other antimitotic agents. Scale bar at bottom right indicates 5 μm. **(E)** Quantification of the percentage of pHH3+/mitotic cells in images from stained tissue sections. Whole tumor sections were imaged, and for each tumor ten images were selected at random for quantification. Plotted is the mean ± SEM where n = 6 to 8 tumors per condition with ≥6724 cells quantified per tumor. The percentage of pHH3+ cells for individual tumors is depicted in grey circles. P values shown were calculated using a Student’s two-tailed t test; n.s. not significant.

Finally, we sought to determine if the synergistic effects observed on *in vivo* PDX tumor growth were related to the synergistic mitotic arrest observed in cultured cells. End-of-study tumors were harvested, fixed, sectioned, and stained with both DAPI and antibodies against pHH3 as a marker of mitotic cells (**Figure 7C**). Immunofluorescence microscopy revealed that treatment with onvansertib and the combination clearly increased the abundance of mitotic cells. Moreover, pHH3 positive staining cells in onvansertib and combination treated tumor sections contained decondensed and dispersed chromatin (**Figure 7D**), a morphology that resembles collapsed/monopolar spindles observed after prolonged Plk1 inhibition (53, 68). We next quantified the percentage of cells that stained positive for pHH3 in tissue sections from all tumors. In line with its mechanism of action, tumors treated with onvansertib had more mitotic cells than vehicle-treated tumors. Tumors treated with both drugs, however, had an even greater percentage of pHH3-positive cells than onvansertib alone. Despite a lack of effect from abiraterone monotherapy, cotreatment resulted in ∼30% increase in pHH3-positive cells compared to onvansertib alone (**Figure 7E**). These CRPC PDX data confirm that abiraterone treatment enhances the antimitotic effects of Plk1 inhibition resulting in synergistic mitotic arrest *in vivo*. Taken together, these organoid and human PDX data demonstrate clear synergistic anti-tumor activity of abiraterone in combination with Plk1 inhibition, and provides strong preclinical support for the ongoing clinical trial (NCT03414034) testing the combination of onvansertib and abiraterone in CRPC patients with metastatic disease.

## DISCUSSION

Our early finding of synergistic CPRC cell death in response to combined abiraterone and Plk1 inhibition, and those of Liu and colleagues, have led to a current clinical investigation (NCT03414034; Einstein et al., 2022) of this drug combination in mCPRC patients (70, 71). However, the mechanism of action of the combination has remain unknown until now. Here we report a cell biological mechanism for synergistic cancer cell killing, demonstrate that this mechanism is independent of androgen receptor signaling and is applicable to cancer types beyond tumors of the prostate. Consistent with our finding of synergistic cell death, Zhang et al showed strong synergy between abiraterone and Plk1 inhibitors in prostate cancer cells, especially with regard to *in vivo* efficacy of combined Plk1 inhibitor volasertib with abiraterone in an AR-v7-positive LuCAP35CR PDX model of CRPC. In that study, however, the authors concluded that the observed synergy was AR-dependent, and resulted from Plk1-dependent effects that markedly potentiated androgen signaling and testosterone synthesis. In contrast, our work, however, shows that this drug combination causes synergistic mitotic arrest and cancer cell death in an AR-independent manner both in culture and *in vivo*, through a process that possibly involves abiraterone-induced defects in spindle orientation and chromatin compaction. The fact that abiraterone-Plk1 inhibitor synergy was observed in two different prostate cancer PDX models (LuCAP35CR used by Zhang *et al.* and LVCaP2-CR shown in **Figure 7**), both of which express the AR-v7 splice variant, further indicates that this synergy is independent of AR signaling since AR-v7 is constitutively active and AR-v7- expressing tumors are generally poorly responsive to abiraterone treatment alone (66, 72). Furthermore, synergistic cell killing by the combination of abiraterone and Plk1 kinase inhibitors was observed in a variety of non-prostate cancer cell lines, none of which had demonstrable expression of the androgen receptor. Finally, our finding that inhibition of Cyp17A1 failed to synergize with Plk1 inhibition for cell killing further indicates that the relevant effects of abiraterone in this regard appear to be independent of its effects on steroid biosynthesis.

The observation that abiraterone has additional effects on cells beyond AR antagonism and Cyp17A1 inhibition is likely due to its resemblance to endogenous steroids, particularly pregnenolone. The additional binding partners of abiraterone that are causal for synergy with Plk1 inhibition may be protein(s) that also bind pregnenolone or similar steroidal molecules. These AR-independent effects of abiraterone likely do not contribute significantly to the efficacy of abiraterone when used as monotherapy, since there is fairly strong cross-resistance between abiraterone and enzalutamide in mCRPC patients. Patients taking enzalutamide after PSA progression on abiraterone had longer progression-free survival than *vice versa*, however, the responses were strongly muted relative to androgen signaling inhibitor-naive patients (73). These data imply that the mechanisms of abiraterone resistance in prostate cancer patients largely overlap with those of enzalutamide (with the *vice versa* being less often the case) and support a shared mechanism of action when abiraterone is used as monotherapy (74). However, the data presented here indicate that these AR-independent effects of abiraterone on mitosis can be leveraged to elicit increased sensitivity to Plk1 inhibition and that this combination would remain useful in the context of antiandrogen resistance, a clinical setting with significant and increasing unmet need (75).

In contrast to the AR-independent effects of abiraterone, Plk1 inhibitors’ role in this synergy were both specific and unique to Plk1 among a panel of antimitotic drugs (**Figure 2I**). Abiraterone sensitized CPRC cells to multiple structurally disparate Plk1 inhibitors synergized with abiraterone, and had no effect on sensitivity to other agents that target processes in mitosis, including docetaxel, nocodazole, Aurora A inhibitors, and Aurora B inhibitions. Thus, it appears there is something unique about the mitotic consequences of Plk1 inhibition that synergize with the androgen-independent effects of abiraterone in some cancer cells.

Notably, we show here that abiraterone treatment prevents anchoring of the mitotic spindle axis with respect to the cell cortex, and impairs complete chromosome condensation, although the precise molecular basis for these effects is not yet known. Both of these processes, which are critical for proper execution of mitotic cell division, are themselves complex and incompletely understood. Spindle orientation and stabilization depends on connections between the astral microtubule network and a complex consisting of membrane-bound Gαi, and the proteins LGN and NuMA at the cell cortex. The Gαi-LGN-NuMA complex accumulates at the cell cortex adjacent to each spindle pole forming a crescent shape. Dynein motors on astral microtubules then interact with NuMA to properly orient and anchor the spindle poles (76). Altered spindle orientation and excessive spindle rotation can result from defects in astral microtubule dynamics or microtubule nucleation at centrosomes, from disruption of the Gαi-LGN-NuMA complex, or from misslocalization of NuMA which is normally regulated by both cell autonomous and extrinsic factors, including the formation of retraction fibers after cell rounding and actin-cytoskeleton-based sensing of extracellular adhesion and mechanosensitive molecules. In addition, there are a numerous other proteins that participate and regulate these processes and have been shown to cause spindle orientation defects when depleted or inhibited (77), any of which could be the target of abiraterone. Similarly, the process of chromosome condensation, despite have been observed for the first time more than a century ago (78), remains only partially understood at the molecular level. In prophase the condensin II complex becomes associated with chromatin and extrudes DNA through its center to produce large loops, which are then further compacted by condensin I following nuclear envelope breakdown and entry into prometaphase. These condensin complexes then associate with each other to form rod-like structures with loops of chromatin emanating radially like a brush (79). Surprisingly, however, depletion of condensins does not lead to chromatin decondensation in most cell types, making it unlikely that these are direct abiraterone targets. Instead, it is thought that multiple additional factors contribute to chromosome condensation including topoisomerase IIα, the kinesin KIF4A, post-translational modification-specific histone interactions, or divalent cations and polyamines that neutralize electrostatic clashes (80, 81), which could potentially be affected by the presence of abiraterone.

Finally, combined abiraterone and Plk1 inhibitor treatment increased the frequency of entotic cell death which was not seen with enzalutamide co-treated cells. Prolonged mitotic arrest has been shown to result in entosis (63) which appears to be the result of the arrested mitotic cells physically invading adjacent neighboring cells. Importantly, we did not observe this phenotype in our previous work using C4-2 cells treated with other drug combinations that cause synergistic mitotic arrest, such as Plk1 inhibitors and microtubule depolymerizing agents (53). Thus, while entosis may be a consequence of mitotic arrest in some circumstances, it is not a general feature of mitotically arrested C4-2 CRPC cells, and its prevalence after combined abiraterone-Plk1 inhibition may implicate particular aspects of mitosis as likely candidates for the mechanism of synergy.

In summary, our data demonstrate a cellular mitotic mechanism rationalizing the use of combined abiraterone and Plk1 inhibitors in patients with mCRPC, and provide a compelling opportunity for repurposing of abiraterone as a novel antimitotic agent, that on its own exerts mild effects on mitosis, but can be combined with Plk1 inhibitors such as onvansertib to amplify these effects and synergistically kill cancer cells. Importantly, our finding of AR-independent effects of abiraterone and significant synergy in non-prostate cancer tumor types provides scientific justification for this combination outside the context of mCRPC. Abiraterone is an exceptionally well-tolerated drug in the field of oncology (8). Enhancing the effects of onvansertib though combination with abiraterone could be an attractive way to more generally enhance the utility of Plk1 inhibitors as anti-cancer agents in a wide variety of different tumor types.

## MATERIALS AND METHODS

### Culturing of Cells

All cell lines were cultured in a 37°C humidified incubator supplied with 5% CO_2_, were maintained subconfluent, and used for no more than 20 passages. All media was supplemented with fetal bovine serum (FBS) unless explicitly stated as charcoal-stripped FBS (csFBS), contained 2 mM glutamine, and lacked antibiotics. C4-2, LNCaP, and PSN-1 were grown in RPMI-1640 with 10% FBS. Panc 10.05 was grown in RPMI-1640 with 15% FBS, and 10 IU/ml human recombinant insulin (Gibco). OAW28 was grown in Advanced DMEM (Gibco) also supplemented with insulin. PC-3 cells were grown in F-K12 media with 10% FBS. DU145 cells were grown in DMEM media with 10% FBS. SK-OV-3 cells were grown in McCoy’s 5A with 10% FBS. MV-4-11 cells were grown in IMDM with 10% FBS. OCI-AML-3 cells were grown in MEM with 20% FBS. All cell lines used in this study were authenticated using short tandem repeat genotyping (Labcorp) and confirmed to be negative for mycoplasma contamination using MycoAlert^TM^ PLUS Assay (Lonza).

### Measurements of Drug Sensitivity and Drug Combination Synergy

Cells were plated in 96- or 384-well plates at a density of 4000 or 1000 cells per well in 22.5 or 95 μl media, respectively. For experiments using the suspension AML cell lines, MV-4-11 and OCI-AML-3, 10,000 cells were plated in 384-well plates in 22.5 μl. The following day drugs, diluted in 5 μl or 2.5 μl media for 96- or 384-well plates, respectively, were added to the wells. A constant amount of DMSO vehicle was maintained in all wells. Unless otherwise stated, after 72-hour incubation, viability relative to control was measured using CellTiter-Glo^TM^ (Promega) according to the manufacturer’s recommendations. Luminescence in individual wells in plates with opaque walls was measured using and Infinite^TM^ M200 Pro plate reader (Tecan Group Ltd.). Relative viability was calculated by dividing the luminescent signal from each well by that measured in the vehicle-control well on an individual replicate basis. All abiraterone used in this study is specifically abiraterone acetate (Selleck Chemicals). For experiments examining cell number and relative confluence, drug treated cells were washed with PBS and fixed with 4% formaldehyde for 1 hour, stained with 1 μM SYTO^TM^ 60 (Molecular Probes) and imaged on an Odyssey^TM^ CLx scanner (LiCOR Biosciences).

### Flow Cytometry Analysis of Cell Cycle Distribution and Apoptosis

The indicated cells were treated as described and then collected by centrifugation or trypsinization for suspension or adherent cells, respectively. The media, trypsin, and PBS wash were collected together to avoid loss of loosely attached or detached cells. Cells were fixed in 4% formaldehyde in PBS for 15 minutes, washed with PBS containing 1% bovine serum albumin (PBS-BSA), and then stored in methanol at -20°C overnight. Cells were then washed twice in PBS-BSA 0.1% Tween-20, incubated with primary antibodies overnight at 4°C, washed with PBS-BSA 0.1% Tween-20, and incubated for 1 hour with fluorescent-dye conjugated secondary antibodies (diluted 1:200, Alexa Fluor, Molecular Probes) at room temperature for 1 hour. The fixed cells were then washed with PBS-BSA 0.1% Tween-20, resuspended in PBS containing 1 μg/ml 4,6-diamidino-2-phenylindole (DAPI, Molecular Probes) to stain DNA, and analyzed using a BD^TM^ LSRII flow cytometer (Becton Dickinson) and the FlowJo^TM^ software package. Primary antibodies used included anti-phospho serine 10 histone H3 (pHH3; EMD Millipore, 3H10) and anti-cleaved capase-3 (BD Pharmingen, 599565). For cell-cycle analysis post Cyp17A1 knockdown, prostate cancer cells were detached by treatment with Accutase (Gibco), fixed in ethanol, resuspended in propidium iodide (PI) staining solution (50 μg/mL PI, 0.1 mg/mL RNase A, and 0.05% Triton X-100), and analyzed on a FACSCanto II (BD Biosciences) as described previously (82).

### Time-lapse Live-cell Microscopy

Live-cell microscopy for analysis of mitotic spindles, duration of mitosis, and phenotypes associated with drug treatment was performed using C4-2 CRPC cells expressing histone H2B-mCherry and mEmerald-tubulin. Cells (125, 000) were plated in 12-well plates using 1.5 mls phenol red free media. The following day drugs were added as indicated and the cells were imaged on a EVOS^TM^ FL Auto Cell Imaging System (Invitrogen) equipped with with an onstage incubator to maintain 37°C, adequate humidity, and 5% CO_2_. Images were acquired at 15-minute intervals for 72 hours using a 20X objective lens. Images were compiled into movies and analyzed using the Fiji distribution of ImageJ v2.1.0 (83). For image analysis of spindle rotation, and cell fate post mitosis, cells to be analyzed per condition were picked at random using only the first frame of the time lapse. Once the cell entered its first mitosis, the angle of the spindle’s major axis was tracked starting from when it was first apparent, prometaphase, until telophase. For each frame transition we calculated both the clockwise and counterclockwise deviations and assumed that the actual rotation was the lesser of the two (the shortest rotational path). Given the modest amount of rotation that occurred in vehicle only treated cells (see **Figure 4E** and **Supplemental Movie 5**), we believe this assumption would rarely, if ever, provide us with the wrong conclusion. Clockwise rotation was considered positive rotation, whereas counterclockwise rotation was negative. The net angular displacement is the total change in angle of the spindle during mitosis from start until finish and is the absolute value of the sum of all stepwise rotations. The cumulative angular distance is the total amount of rotation regardless of direction that occurred during mitosis and is the sum of the absolute value of all rotations.

### Indirect Immunofluorescence of fixed cells and tumor sections

For examining mitotic spindles in fixed cells, C4-2 cells were plated in 12-well plates containing poly-L-lysine coated glass coverslips (Corning-BioCoat #354085) and subjected to the indicated drugs the following day. After the indicated amount of time cells were washed for 1 minute with microtubule stabilization buffer (MTSB: 4 M glycerol, 100 mM PIPES pH 6.9, 1 mM EGTA, 5 mM MgCl_2_), 2 minutes with MTSB 0.5% Triton X-100, 2 minutes with MTSB, 5 minutes with calcium buffer (100 mM PIPES, pH 6.8, 1 mM MgCl_2_, 1 mM CaCl_2_, 0.5% Triton X-100) and then fixed for 10 minutes with 1% glutaraldehyde (Electron Microscopy Sciences) in PBS. Residual glutaraldehyde was quenched with NaBH_4_ (0.5 mg/ml in ddH2O) for 12 minutes twice and then washed twice with TBS-BSA (10 mM Tris-HCl pH 7.5, 150 mM NaCl, 5% BSA) for 5 minutes. Fixed cells were blocked for 1 hour in TBS-BSA containing 0.5% Triton X-100. TBS-BSA was used for anti-α-Tubulin (Novus Biologicals, YL1/2) and anti-CENP-A (Thermo Fisher, PA5-17194) antibody incubation overnight at 4°C, followed by a 5-minute wash, fluorescent secondary antibody (1:200, Alexa Fluor, Molecular Probes) incubation 1 hour at room temperature, and an additional 5-minute wash. Cells were stained using 5 mg/ml DAPI, washed in PBS for 3 minutes, and coverslips mounted on glass slides using the antifade reagent ProLong Gold ^TM^ (Molecular Probes). Images were acquired using a Deltavision ^TM^ Ultimate Focus microscope (Applied Precision) using a 100X 1.40 NA objective. Z-stacks were deconvolved and maximum intensity Z-stack projections were generated using softWoRx ^TM^ software (Applied Precision). The Fiji distribution of ImageJ was used for image analysis. For immunofluorescence analysis of DU145 cancer cells, 1 – 1.5 × 10^4^ were seeded in each well of a 4-chamber slide in 500 mL of culture medium and treated as indicated for 12 hours. Cell culture medium was removed and cells were rinsed with PBS three times. Cell fixation was done using 400 μL of 4% paraformaldehyde (pH 7.4) for 10 min at 37°C followed by 400 μL of ice-cold methanol for 5 min at 20°C. Fixed cells were permeabilized with 0.1% Triton X-100 in PBS (room temperature for 15 min), and blocking was done with 2% BSA in PBS (room temperature for 60 min). The anti-α-Tubulin primary antibodies incubation was performed overnight. Fluorescent dye–labeled secondary antibody along with DAPI diluted in 500 μL of 0.1% BSA was added for 45 min in room temperature. Negative and positive controls were used. Z-stacks were acquired and deconvolved as above but using a 60X 1.4 NA objective. The analysis using ImageJ was done by two independent investigators.

For immunohistochemical staining of tumor samples, formalin-fixed paraffin-embedded tumors were sectioned and mounted on charged slides. Sections (4 μm) were quenched with 3% H_2_O_2_ (UltraVision Hydrogen Peroxide Block, Epredia), blocked with Rodent Block M (BioCare Medical), labeled with anti-phospho histone H3 antibody (pHH3; Cell Signaling Technology, 9701S) and an Alexa Fluor® 488 conjugated secondary antibody (Molecular Probes), and then stained with DAPI. Images were acquired using a Deltavision ^TM^ Ultimate Focus microscope using a 40X 0.85 NA objective. Randomly selected fields of cells were analyzed using QuPath v0.2.3 (84). Images that contained very few cells or large necrotic regions were replaced with a new randomly selected image. Cells were identified in DAPI images and then pHH3 positive cells were quantified. Automated cell identification and pHH3 quantification was manually inspected for accuracy to ensure accurate cell counting and avoidance of false positive and negative pHH3 assignment to cells.

### Preparation of Lysates and Immunoblotting

Cells were lysed directly on the plate after washing with PBS using lysis buffer (50 mM Tris-HCl pH 6.8, 2% SDS, 5% glycerol, 5 mM EDTA, 1 mM NaF, 10 mM β-glycerophosphate, 1 mM phenylmethylsulfonyl fluoride, 1 mM Na_3_VO_4_, cOmplete^TM^ EDTA-free protease inhibitors, and PhosSTOP^TM^ phosphatase inhibitors). The media and PBS wash were reserved, centrifuged, and any cells present were combined with the lysate to prevent loss of loose or unattached cells. After sonication and protein concentration normalization, 6X sodium dodecyl sulfate - polyacrylamide gel electrophoresis (SDS-PAGE) loading buffer (208 mM Tris-HCl pH 6.8, 42% glycerol, 3 M β-mercaptoethanol, 10% SDS, 5 mg/ml bromophenol blue) was added and lysates were boiled for 5 minutes. Following SDS-PAGE, immunoblots were blocked with Odyssey^TM^ blocking buffer (LiCOR Biosciences) and incubated with primary antibody overnight at 4°C and then secondary antibody for 1 hour at room temperature. Immunoblots were scanned on a Odyssey^TM^ CLx scanner (LiCOR Biosciences). Antibodies used included anti-AR (Cell Signaling Technologies, D6F11), anti-MAD2 (Bethyl Labs, A300-300A), anti-Cyp17A1 (Cell Signaling Technologies, E6Y3S), anti-Vinculin (Santa Cruz Biotechnology, 7F9), and anti-β-actin (Sigma Aldrich, AC-15).

### siRNA Knockdown and Drug Sensitivity Analysis

*Silencer*^®^ Select siRNAs (s8392, Invitrogen) targeting Mad2 (gene name *MAD2L1*) and the Negative Control No. 1 siRNA (#4390844, Invitrogen) were used for Lipofectamine^TM^ RNAiMAX (Thermo Fisher Scientific) based transfection at 10 nM concentration according to the manufacturer’s recommendations. After 24 hours, cells were replated into both 6- and 384-well plates for preparation of cell lysates and synergy experiments, respectively. The following day cells in the 6-well plates were lysed for immunoblots, whereas cells in the 384-well plates were treated with a combination drug matrix that was assessed for synergy after 72 hours. For siCYP17A1 experiments Dharmacon On-TARGETplus ^TM^ human CYP17A1 siRNA (LQ-008469-02-0005) were used.

### Gene Expression Analysis by RNA Sequencing

Cells were plated in 6-well plates (300,000 cells per well), and 24 hours later treated with the indicated drugs in biological triplicate. After 16 hours cells were lysed and total RNA was isolated using NucleoSpin^®^ RNA Plus Mini Kit (Macherey-Nagel) according to the manufacturer’s recommendations. Samples were submitted to the MIT BioMicro Center for library preparation and sequencing. RNA quality was assessed using a Fragment Analyzer (Agilent Technologies) and RNA sequencing libraries were prepared using 400 ng of total RNA using the Kapa mRNA Hyperprep kit (Roche) at 1/3rd reaction volume using 14 cycles of PCR. Libraries were analyzed using the Fragment Analyzer and quantified by qPCR prior to pooling and sequencing on a NextSeq^®^ 500 System (Illumina) using 75nt single end reads. Sequences were aligned to the human transcriptome (Gencode v29 GRCh38.p12)followed by transcript quantification using Salmon v0.14 (85). Transcript abundance was transformed into gene set enrichment scores using GSVA v1.36 (57) and MSigDB v7.0, the relevant contrasts performed using limma v3.44 (58), and differentially expressed gene sets identified based on an ≤ 0.01 FDR cut off. Area-proportional Venn diagrams were made using EulerAPE v3 (86).

### Organoid Generation, Culture, and Drug Sensitivity Measurements

Prostate cancer 3D organoids were derived from PDXs grown in castrated male NOD scid mice (Taconic). Following euthanasia tumors were extracted. Minced tumor fragments for organoid culturing were digested in Accumax^TM^ - Cell Aggregate Dissociation Medium (Thermo Fisher Scientific), resuspended in DMEM/F12 plus 10% FBS and passed through a 250-µm cell strainer (Thermo Fisher Scientific, Pierce Tissue Strainers, #87791) to remove tissue debris and obtain smaller cell clusters. Cells were plated on Matrigel covered tissue cultured plates (Corning Matrigel^®^, Growth Factor Reduced Basement Membrane Matrix, LDEV-free, # 354230) and prostate-organoid specific media (87). After 7 days, organoids were transferred to 96-well plates and treated with onvansertib, abiraterone, and the combination for 6 days. Cell viability was assessed with CellTiter-Glo^®^ 3D Cell Viability Assay (Promega). Six biological replicates were assessed per treatment group.

### In vivo Studies Using a Tumor-Implantable Microwell Device

C4-2 CRPC and SK-OV-3 ovarian cancer xenograft tumors were grown in four- to six-week old NCR nude mice (Taconic), male or female, respectively. Five million C4-2 or two million SK-OV-3 cells in serum free media were mixed one to one with growth factor reduced Matrigel^®^ (Invitrogen) in a total volume of 200 μl and injected in the hind flank using a 23- or 27-gauge needle, respectively. Cells were found to be free of murine pathogens by IMPACT rodent pathogen testing (IDEXX BioAnalytics) prior to injection. Tumors took four to eight weeks to grow.

Microdose containing tumor-implantable devices were manufactured, implanted, and *in vivo* drug responses analyzed as previously described (65). The cylindrical microdevices (4 mm x 820 μm) micromachined from medical-grade Delrin^®^ acetal resin blocks (DuPont) each contain eighteen 200 µm (diameter) x 250 µm (depth) reservoirs. Abiraterone acetate and BI2536 were mixed with PEG 1450 at 12.5% by weight, and 1 µg of the dry powder mixture then packed into a reservoir. Wells containing the combination of drugs contained 12.5% of each drug by weight. Implantation of the devices was accomplished using a 19-gauge spinal biopsy needle (Angiotech) with a retractable needle obturator. Tumors were excised 24 to 36 hours after device implantation, fixed in 10% formalin for 24 hours, and embedded in paraffin. Sections were stained with cleaved caspase-3 antibody (Cell Signaling Technologies, 9664) followed by detection with horseradish peroxidase conjugated secondary antibody and diaminobenzidine with hematoxylin used as a counterstain, following standard immunohistochemistry techniques. Images were acquired using an EVOS^®^ Cell Imaging System (Invitrogen) microscope, and scored using ImageJ (83) in a blinded manner. If upon implantation a well was adjacent to necrotic tissue then the response to the drugs could not be assessed. Because of this the number of measurements is not the same between treatment groups and ranged from 5 to 13. The apoptotic index was calculated as the percentage of cells that stained positive for cleaved caspase-3 within a 400 μm radius of the microwell-tissue interface, as described in (88).

All animal studies were approved by the Massachusetts Institute of Technology Committee for Animal Care or Beth Israel Deaconess Institutional Animal Care and Use Committee, conducted in compliance with both the Animal Welfare Act Regulations and other federal statutes relating to animals and experiments involving animals, and adhered to the principles set forth in the Guide for the Care and Use of Laboratory Animals, National Research Council, 1996 (Institutional Animal Welfare Assurance #A-3125–01).

### In vivo Studies Using LVCaP-2CR PDX

LVCaP-2CR PDXs were grown in castrated male NOD scid gamma mice obtained from an in-house colony at the Sidney Kimmel Comprehensive Cancer Center (SKCCC) at Johns Hopkins as described previously (66). Twenty milligrams of LVCaP-2CR contained in 200 µl of 50% Matrigel^®^ (Invitrogen) was subcutaneously injected in the hind flank of the mice. After 4 weeks, 0.1 cm^3^ tumors had formed and the animals were randomized into 4 groups. Onvansertib was suspended in 0.5% methylcellulose 0.1% Tween 80 and administered at 60 mg/kg. Abiraterone acetate (Selleck Chem) was administered at 0.5 mmole/kg (i.e. 196 mg/kg) in 200 μl of 5% benzl alcohol 95% safflower oil. Both were administered by oral gavage. Animals were treated in cycles consisting of 5 consecutive days on treatment, followed by a 24 to 48-hour drug holiday. Tumor size was measured at the indicated days and volume was calculated according to the ellipsoid volume formula (length × width × height × π/6). All animal procedures for the LVCaP-2CR PDX study were approved by the Johns Hopkins University School of Medicine Institutional Animal Care and Use Committee.

## ACKNOWLEDGEMENTS

We thank past and present members of the Yaffe lab, particularly Drs. Yi-Wen Kong, Daniel Lim, and Brian Joughin, and Ian Hickson (Newcastle University) for helpful advice and discussions. We acknowledge the Koch Institute Genomics Core and MIT BioMicro Center and the Koch Institute’s Robert A. Swanson (1969) Biotechnology Center for technical support, specifically the Flow Cytometry Core Facility, The Microscopy Core Facility, and the Hope Babette Tang (1983) Histology Facility. This work was supported in part by the Koch Institute Support (core) Grant P30-CA14051 from the National Cancer Institute. We wish to thank the SKCCC at Hopkins Animal Core Facility supported by the SKCCC CCSG (P30 CA006973) for their services and assistance. This research was supported by grants from the National Institutes of Health (R35-ES028374 to M.B.Y.; P01 CA163227, and P01 P50 CA090381 to S.P.B.), awards from the Charles and Marjorie Holloway Foundation, the Ovarian Cancer Research Fund, the Prostate Cancer Foundation and the MIT Center for Precision Cancer Medicine to M.B.Y., two Koch Institute - Dana-Farber/Harvard Cancer Center Bridge Project Grants to M.B.Y., S.P.B., and D.J.E., an American Cancer Society Post-Doctoral Fellowship PF-13-355-01-TBE to J.C.P., a Prostate Cancer Foundation Young Investigator Award to D.J.E., the Janssen/TRANSCEND Program, and a Sponsored Research Agreement from Cardiff Oncology Inc to M.B.Y.

## Notes

**DISCLOSURE OF POTENTIAL CONFLICT OF INTERESTS** Both Michael B. Yaffe and Jesse C. Patterson are co-inventors on a patent for the use of antiandrogens in combination with inhibitors of Plk1 (U.S. Patent and Trademark Office: 9,566,280). That patent has been licensed by Cardiff Oncology Inc. that owns the rights to onvansertib. Peter J.P. Croucher, Maya Ridinger, and Mark G. Erlander are employees and shareholders of Cardiff Oncology, Inc.

### Competing Interest Statement

Both Michael B. Yaffe and Jesse C. Patterson are co-inventors on a patent for the use of antiandrogens in combination with inhibitors of Plk1 (U.S. Patent and Trademark Office: 9,566,280). That patent has been licensed by Cardiff Oncology Inc. that owns the rights to onvansertib. Peter J.P. Croucher, Maya Ridinger, and Mark G. Erlander are employees and shareholders of Cardiff Oncology, Inc.

## REFERENCES

1. Augello MA, Den RB, Knudsen KE. AR function in promoting metastatic prostate cancer. Cancer and Metastasis Reviews 2014;33:399–411.

2. Yuan X, Cai C, Chen S, Chen S, Yu Z, Balk SP. Androgen receptor functions in castration-resistant prostate cancer and mechanisms of resistance to new agents targeting the androgen axis. Oncogene 2014;33:2815–25.

3. Attard G, Belldegrun AS, de Bono JS. Selective blockade of androgenic steroid synthesis by novel lyase inhibitors as a therapeutic strategy for treating metastatic prostate cancer. BJU International 2005;96:1241–6.

4. Cai C, Chen S, Ng P, Bubley GJ, Nelson PS, Mostaghel E a, et al. Intratumoral de novo steroid synthesis activates androgen receptor in castration-resistant prostate cancer and is upregulated by treatment with CYP17A1 inhibitors. Cancer Res 2011;71:6503–13.

5. Li Z, Bishop AC, Alyamani M, Garcia JA, Dreicer R, Bunch D, et al. Conversion of abiraterone to D4A drives anti-tumour activity in prostate cancer. Nature 2015;523:347–51.

6. Richards J, Lim AC, Hay CW, Taylor AE, Wingate A, Nowakowska K, et al. Interactions of abiraterone, eplerenone, and prednisolone with wild-type and mutant androgen receptor: A rationale for increasing abiraterone exposure or combining with MDV3100. Cancer Research 2012;72:2176–82.

7. Jung ME, Ouk S, Yoo D, Sawyers CL, Chen C, Tran C, et al. Structure-activity relationship for thiohydantoin androgen receptor antagonists for castration-resistant prostate cancer (CRPC). Journal of Medicinal Chemistry 2010;53:2779–96.

8. Ryan CJ, Smith MR, de Bono JS, Molina A, Logothetis CJ, de Souza P, et al. Abiraterone in metastatic prostate cancer without previous chemotherapy. New England Journal of Medicine 2013;368:138–48.

9. Beer TM, Armstrong AJ, Rathkopf DE, Loriot Y, Sternberg CN, Higano CS, et al. Enzalutamide in metastatic prostate cancer before chemotherapy. New England Journal of Medicine 2014;371:424–33.

10. Su Z, Zhang M, Xu M, Li X, Tan J, Xu Y, et al. MicroRNA181c inhibits prostate cancer cell growth and invasion by targeting multiple ERK signaling pathway components. Prostate 2018;78:343–52.

11. Carver BS, Chapinski C, Wongvipat J, Hieronymus H, Chen Y, Chandarlapaty S, et al. Reciprocal feedback regulation of PI3K and androgen receptor signaling in PTEN-deficient prostate cancer. Cancer Cell 2011;19:575–86.

12. Culig Z, Hobisch A, Cronauer MV, Radmayr C, Trapman J, Hittmair A, et al. Androgen receptor activation in prostatic tumor cell lines by insulin-like growth factor-1, keratinocyte growth factor and epidermal growth factor. European Urology 1995;27:45–7.

13. Yardy GW, Brewster SF. Wnt signalling and prostate cancer. Prostate Cancer and Prostatic Diseases 2005;8:119–26.

14. Ferraldeschi R, Nava Rodrigues D, Riisnaes R, Miranda S, Figueiredo I, Rescigno P, et al. PTEN protein loss and clinical outcome from castration-resistant prostate cancer treated with abiraterone acetate. European Urology 2015;67:795–802.

15. Miyamoto DT, Zheng Y, Wittner BS, Lee RJ, Zhu H, Broderick KT, et al. RNA-Seq of single prostate CTCs implicates noncanonical Wnt signaling in antiandrogen resistance. Science 2015;349:1351–6.

16. Deeraksa A, Pan J, Sha Y, Liu X-D, Eissa NT, Lin S-H, et al. Plk1 is upregulated in androgen-insensitive prostate cancer cells and its inhibition leads to necroptosis. Oncogene 2013;32:2973– 83.

17. Liu XS, Song B, Elzey BD, Ratliff TL, Konieczny SF, Cheng L, et al. Polo-like kinase 1 facilitates loss of Pten tumor suppressor-induced prostate cancer formation. J Biol Chem 2011;286:35795– 800.

18. Zhang W, van Gent DC, Incrocci L, van Weerden WM, Nonnekens J. Role of the DNA damage response in prostate cancer formation, progression and treatment. Prostate Cancer and Prostatic Diseases 2020;23:24–37.

19. Oh M, Alkhushaym N, Fallatah S, Althagafi A, Aljadeed R, Alsowaida Y, et al. The association of BRCA1 and BRCA2 mutations with prostate cancer risk, frequency, and mortality: A meta-analysis. Prostate 2019;79:880–95.

20. Gallagher DJ, Gaudet MM, Pal P, Kirchhoff T, Balistreri L, Vora K, et al. Germline BRCA mutations denote a clinicopathologic subset of prostate cancer. Clinical Cancer Research 2010;16:2115–21.

21. Cybulski C, Wokołorczyk D, Kluźniak W, Jakubowska A, Górski B, Gronwald J, et al. An inherited NBN mutation is associated with poor prognosis prostate cancer. British Journal of Cancer 2013;108:461–8.

22. Grindedal EM, Møller P, Eeles R, Stormorken AT, Bowitz-Lothe IM, Landrø SM, et al. Germ-line mutations in mismatch repair genes associated with prostate cancer. Cancer Epidemiology Biomarkers and Prevention 2009;18:2460–7.

23. Dominguez-Valentin M, Joost P, Therkildsen C, Jonsson M, Rambech E, Nilbert M. Frequent mismatch-repair defects link prostate cancer to Lynch syndrome. BMC Urology 2016;16:1–7.

24. Fraser M, Sabelnykova VY, Yamaguchi TN, Heisler LE, Livingstone J, Huang V, et al. Genomic hallmarks of localized, non-indolent prostate cancer. Nature 2017;541:359–64.

25. Potugari BR, Engel JM, Onitilo AA. Metastatic prostate cancer in a RAD51C mutation carrier. Clinical Medicine and Research 2018;16:69–72.

26. Na R, Zheng SL, Han M, Yu H, Jiang D, Shah S, et al. Germline Mutations in ATM and BRCA1/2 Distinguish Risk for Lethal and Indolent Prostate Cancer and are Associated with Early Age at Death. European Urology 2017;71:740–7.

27. Kaur H, Salles DC, Murali S, Hicks JL, Nguyen M, Pritchard CC, et al. Genomic and Clinicopathologic Characterization of ATM-deficient Prostate Cancer. Clinical Cancer Research 2020;26:4869–81.

28. Robinson D, van Allen EM, Wu YM, Schultz N, Lonigro RJ, Mosquera JM, et al. Integrative clinical genomics of advanced prostate cancer. Cell 2015;161:1215–28.

29. Wokołorczyk D, Kluźniak W, Stempa K, Rusak B, Huzarski T, Gronwald J, et al. PALB2 mutations and prostate cancer risk and survival. British Journal of Cancer 2021;125:569–75.

30. Holder J, Poser E, Barr FA. Getting out of mitosis: spatial and temporal control of mitotic exit and cytokinesis by PP1 and PP2A. FEBS Letters 2019;593:2908–24.

31. Archambault V, Glover DM. Polo-like kinases: Conservation and divergence in their functions and regulation. Nature Reviews Molecular Cell Biology 2009;10:265–75.

32. Takai N, Hamanaka R, Yoshimatsu J, Miyakawa I. Polo-like kinases (Plks) and cancer. Oncogene 2005;24:287–91.

33. Weichert W, Schmidt M, Gekeler V, Denkert C, Stephan C, Jung K, et al. Polo-like kinase 1 is overexpressed in prostate cancer and linked to higher tumor grades. Prostate 2004;60:240–5.

34. Witkiewicz AK, Chung S, Brough R, Vail P, Franco J, Lord CJ, et al. Targeting the Vulnerability of RB Tumor Suppressor Loss in Triple-Negative Breast Cancer. Cell Reports 2018;22:1185–99.

35. Degenhardt Y, Greshock J, Laquerre S, Gilmartin AG, Jing J, Richter M, et al. Sensitivity of Cancer Cells to Plk1 Inhibitor GSK461364A Is Associated with Loss of p53 Function and Chromosome Instability. Molecular Cancer Therapeutics 2010;9:2079–89.

36. Wu J, Ivanov AI, Fisher PB, Fu Z. Polo-like kinase 1 induces epithelial-to-mesenchymal transition and promotes epithelial cell motility by activating CRAF/ERK signaling. Elife 2016;5:1–25.

37. Gheghiani L, Shang S, Fu Z. Targeting the PLK1-FOXO1 pathway as a novel therapeutic approach for treating advanced prostate cancer. Scientific Reports 2020;10:1–11.

38. Cristóbal I, Rojo F, Madoz-Gúrpide J, García-Foncillas J. Cross Talk between Wnt/β-Catenin and CIP2A/Plk1 Signaling in Prostate Cancer: Promising Therapeutic Implications. Molecular and Cellular Biology 2016;36:1734–9.

39. Lehár J, Krueger AS, Avery W, Heilbut AM, Johansen LM, Price ER, et al. Synergistic drug combinations tend to improve therapeutically relevant selectivity. Nat Biotechnol 2009;27:659– 66.

40. Lorente D, Mateo J, Perez-Lopez R, de Bono JS, Attard G. Sequencing of agents in castration-resistant prostate cancer. The Lancet Oncology 2015;16:e279–92.

41. Li K, Zhan W, Chen Y, Jha RK, Chen X. Docetaxel and doxorubicin codelivery by nanocarriers for synergistic treatment of prostate cancer. Frontiers in Pharmacology 2019;10:1–16.

42. Polkinghorn WR, Parker JS, Lee MX, Kass EM, Spratt DE, Iaquinta PJ, et al. Androgen receptor signaling regulates DNA repair in prostate cancers. Cancer Discov 2013;3:1245–53.

43. Kishi K, van Vugt MATM, Okamoto K, Hayashi Y, Yaffe MB. Functional Dynamics of Polo-Like Kinase 1 at the Centrosome. Molecular and Cellular Biology 2009;29:3134–50.

44. Vugt MATM van, Gardino AK, Linding R, Ostheimer GJ, Reinhardt C, Ong S, et al. A Mitotic Phosphorylation Feedback Network Connects Cdk1, Plk1, 53BP1, and Chk2 to Inactivate the G2/M DNA Damage Checkpoint. PLoS Biol 2010;8:e1000287.

45. Macůrek L, Lindqvist A, Lim D, Lampson MA, Klompmaker R, Freire R, et al. Polo-like kinase-1 is activated by aurora A to promote checkpoint recovery. Nature 2008;455:119–23.

46. Elia AEH, Rellos P, Haire LF, Chao JW, Ivins FJ, Hoepker K, et al. The molecular basis for phosphodependent substrate targeting and regulation of Plks by the Polo-box domain. Cell 2003;115:83–95.

47. Elia AEH, Cantley LC, Yaffe MB. Proteomic screen finds pSer/pThr-binding domain localizing Plk1 to mitotic substrates. Science 2003;299:1228–31.

48. Lowery DM, Clauser KR, Hjerrild M, Lim D, Alexander J, Kishi K, et al. Proteomic screen defines the Polo-box domain interactome and identifies Rock2 as a Plk1 substrate. EMBO Journal 2007;26:2262–73.

49. Wang Q, Li W, Zhang Y, Yuan X, Xu K, Yu J, et al. Androgen receptor regulates a distinct transcription program in androgen-independent prostate cancer. Cell 2009;138:245–56.

50. Hu R, Lu C, Mostaghel EA, Yegnasubramanian S, Gurel M, Tannahill C, et al. Distinct transcriptional programs mediated by the ligand-dependent full-length androgen receptor and its splice variants in castration-resistant prostate cancer. Cancer Research 2012;72:3457–62.

51. Lee MJ, Ye AS, Gardino AK, Heijink AM, Sorger PK, MacBeath G, et al. Sequential application of anticancer drugs enhances cell death by rewiring apoptotic signaling networks. Cell 2012;149:780–94.

52. Bliss CI. THE TOXICITY OF POISONS APPLIED JOINTLY. Annals of Applied Biology 1939;26:585–615.

53. Patterson JC, Joughin BA, Prota AE, Mühlethaler T, Jonas OH, Whitman MA, et al. VISAGE Reveals a Targetable Mitotic Spindle Vulnerability in Cancer Cells. Cell Systems 2019;9:74–92.e8.

54. Wu H -C, Hsieh J -T, Gleave ME, Brown NM, Pathak S, Chung LWK. Derivation of androgen-independent human LNCaP prostatic cancer cell sublines: Role of bone stromal cells. International Journal of Cancer 1994;57:406–12.

55. Gutteridge REA, Ndiaye MA, Liu X, Ahmad N. Plk1 inhibitors in cancer therapy: From laboratory to clinics. Molecular Cancer Therapeutics 2016;15:1427–35.

56. Nguyen HM, Vessella RL, Morrissey C, Brown LG, Coleman IM, Higano CS, et al. LuCaP Prostate Cancer Patient-Derived Xenografts Reflect the Molecular Heterogeneity of Advanced Disease and Serve as Models for Evaluating Cancer Therapeutics. Prostate 2017;77:654–71.

57. Hänzelmann S, Castelo R, Guinney J. GSVA: Gene set variation analysis for microarray and RNA-Seq data. BMC Bioinformatics 2013;14:7.

58. Ritchie ME, Phipson B, Wu D, Hu Y, Law CW, Shi W, et al. Limma powers differential expression analyses for RNA-sequencing and microarray studies. Nucleic Acids Research 2015;43:e47.

59. Liberzon A, Birger C, Thorvaldsdóttir H, Ghandi M, Mesirov JP, Tamayo P. The Molecular Signatures Database Hallmark Gene Set Collection. Cell Systems 2015;1:417–25.

60. Samejima K, Samejima I, Vagnarelli P, Ogawa H, Vargiu G, Kelly DA, et al. Mitotic chromosomes are compacted laterally by KIF4 and condensin and axially by topoisomerase IIα. Journal of Cell Biology 2012;199:755–70.

61. Samoshkin A, Arnaoutov A, Jansen LET, Ouspenski I, Dye L, Karpova T, et al. Human condensin function is essential for centromeric chromatin assembly and proper sister kinetochore orientation. PLoS ONE 2009;4:e6831.

62. Galluzzi L, Vitale I, Aaronson SA, Abrams JM, Adam D, Agostinis P, et al. Molecular mechanisms of cell death: Recommendations of the Nomenclature Committee on Cell Death 2018. Cell Death and Differentiation 2018;25:486–541.

63. Durgan J, Tseng YY, Hamann JC, Domart MC, Collinson L, Hall A, et al. Mitosis can drive cell cannibalism through entosis. Elife 2017;6:1–26.

64. Musacchio A. The Molecular Biology of Spindle Assembly Checkpoint Signaling Dynamics. Current Biology 2015;25:R1002–18.

65. Jonas O, Landry HM, Fuller JE, Santini JT, Baselga J, Tepper RI, et al. An implantable microdevice to perform high-throughput in vivo drug sensitivity testing in tumors. Sci Transl Med 2015;7:284ra57.

66. Zhu Y, Dalrymple SL, Coleman I, Zheng SL, Xu J, Hooper JE, et al. Role of androgen receptor splice variant-7 (AR-V7) in prostate cancer resistance to 2nd-generation androgen receptor signaling inhibitors. Oncogene 2020;39:6935–49.

67. Wu J, Tracey L, Davidoff AM. Assessing interactions for fixed-dose drug combinations in tumor xenograft studies. Journal of Biopharmaceutical Statistics 2012;22:535–43.

68. Steegmaier M, Hoffmann M, Baum A, Garin-chesa P, Lieb S, Adolf R, et al. BI 2536, a Potent and Selective Inhibitor of Polo-like Kinase 1, Inhibits Tumor Growth In Vivo. Current Biology 2007;316–22.

69. Einstein DJ, Choudhury AD, Saylor PJ, Patterson JC, Croucher P, Ridinger M, et al. A phase 2 study of onvansertib in combination with abiraterone and prednisone in patients with metastatic castration-resistant prostate cancer (mCRPC). Journal of Clinical Oncology 2022;40:TPS219– TPS219.

70. Yaffe M, Patterson J. Combination therapies and methods of use thereof for treating cancer. United States Patent Office; 2014. 9,566,280.

71. Zhang Z, Hou X, Shao C, Li J, Cheng JX, Kuang S, et al. PIk1 inhibition enhances the efficacy of androgen signaling blockade in castration-resistant prostate cancer. Cancer Research 2014;74:6635–47.

72. Antonarakis ES, Lu C, Wang H, Luber B, Nakazawa M, Roeser JC, et al. AR-V7 and resistance to enzalutamide and abiraterone in prostate cancer. New England Journal of Medicine 2014;371:1028–38.

73. Khalaf DJ, Annala M, Taavitsainen S, Finch DL, Oja C, Vergidis J, et al. Optimal sequencing of enzalutamide and abiraterone acetate plus prednisone in metastatic castration-resistant prostate cancer: a multicentre, randomised, open-label, phase 2, crossover trial. The Lancet Oncology 2019;20:1730–9.

74. Buck SAJ, Koolen SLW, Mathijssen RHJ, de Wit R, van Soest RJ. Cross-resistance and drug sequence in prostate cancer. Drug Resistance Updates 2021;56:100761.

75. Fizazi K, Tran N, Fein L, Matsubara N, Rodriguez-Antolin A, Alekseev BY, et al. Abiraterone plus Prednisone in Metastatic, Castration-Sensitive Prostate Cancer. New England Journal of Medicine 2017;377:352–60.

76. Bergstralh DT, Dawney NS, St Johnston D. Spindle orientation: A question of complex positioning. Development 2017;144:1137–45.

77. Lechler T, Mapelli M. Spindle positioning and its impact on vertebrate tissue architecture and cell fate. Nature Reviews Molecular Cell Biology 2021;22:691–708.

78. Flemming W. Zellsubstanz, Kern und Zelltheilung. F.C.W. Vogel; 1892.

79. Davidson IF, Peters JM. Genome folding through loop extrusion by SMC complexes. Nature Reviews Molecular Cell Biology 2021;22:445–64.

80. Antonin W, Neumann H. Chromosome condensation and decondensation during mitosis. Current Opinion in Cell Biology 2016;40:15–22.

81. Moser SC, Swedlow JR. How to be a mitotic chromosome. Chromosome Research 2011;19:307– 19.

82. Nouri M, Caradec J, Lubik AA, Li N, Hollier BG, Takhar M, et al. Therapy-induced developmental reprogramming of prostate cancer cells and acquired therapy resistance. Oncotarget 2017;8:18949–67.

83. Schindelin J, Arganda-Carreras I, Frise E, Kaynig V, Longair M, Pietzsch T, et al. Fiji: An open-source platform for biological-image analysis. Nature Methods 2012;9:676–82.

84. Bankhead P, Loughrey MB, Fernández JA, Dombrowski Y, McArt DG, Dunne PD, et al. QuPath: Open source software for digital pathology image analysis. Scientific Reports 2017;7:1–7.

85. Patro R, Duggal G, Love MI, Irizarry RA, Kingsford C. Salmon provides fast and bias-aware quantification of transcript expression. Nature Methods 2017;14:417–9.

86. Micallef L, Rodgers P. euler APE: Drawing area-proportional 3-Venn diagrams using ellipses. PLoS ONE 2014;9:e101717.

87. Puca L, Bareja R, Prandi D, Shaw R, Benelli M, Karthaus WR, et al. Patient derived organoids to model rare prostate cancer phenotypes. Nature Communications 2018;9:1–10.

88. Ahn SW, Ferland B, Jonas OH. An Interactive Pipeline for Quantitative Histopathological Analysis of Spatially Defined Drug Effects in Tumors. J Pathol Inform 2021;12:34.

89. Subramanian A, Tamayo P, Mootha VK, Mukherjee S, Ebert BL, Gillette MA, et al. Gene set enrichment analysis: A knowledge-based approach for interpreting genome-wide expression profiles. Proc Natl Acad Sci U S A 2005;102:15545–50.

