## Supplemental Figures 1 to 5 for "Polo-like kinase-1 Inhibitors and the Antiandrogen Abiraterone Synergistically Disrupt Mitosis and Kill Cancer Cells of Disparate Origin Independently of Androgen Receptor Signaling"

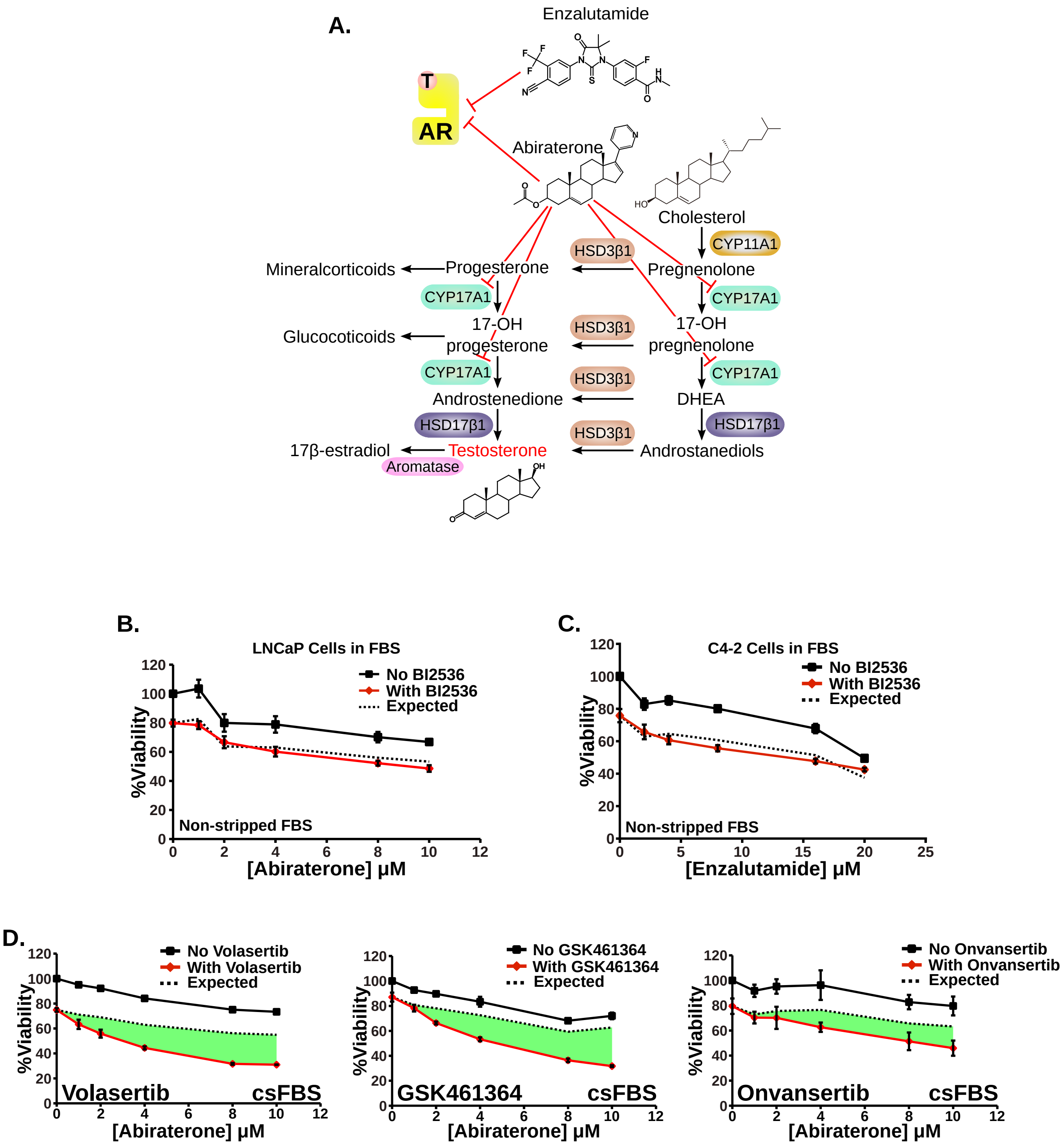


SUPPLEMENTARY FIGURE 1 - Androgen signaling, antiandrogens and synergy between Plk1 inhibition and abiraterone

**Supplemental Figure 1. Androgen signaling, antiandrogens, and synergy between Plk1 inhibition and abiraterone.**

**(A)** Schematic of steroidogenesis and antiandrogen mechanism of action. Abiraterone inhibits Cyp17A1, an enzyme that catalyzes multiple steps in the conversion of cholesterol to testosterone. Both abiraterone and enzalutamide can directly inhibit the AR.

**(B)** LNCaP androgen-dependent cells were grown in non-stripped serum to examine synergy between the Plk1 inhibitor BI2536 (5 nM) and abiraterone. Viability relative to control at 72 hours was measured and mean ± SEM (n = 3) is shown. Expected viability (dotted black line) according to the Bliss Independence model of drug additivity is plotted for comparison. Similar to **Figure 1J** (cells grown in csFBS). No synergy was seen in LNCaP cells in either medium.

**(C)** C4-2 CRPC cells were grown in media containing non-charcoal-stripped FBS and subjected to increasing concentrations of enzalutamide in the absence or presence of the Plk1 inhibitor BI2536 (2.5 nM). Synergy was assessed, analyzed and plotted as in (B). Similar to **Figure 2C** (cells grown in csFBS), enzalutamide and Plk1 inhibitors do not synergistically kill C4-2 CPRC cells when grown in normal FBS.

**(D)** C4-2 CRPC cells were grown in media containing charcoal-stripped FBS and subjected to increasing concentrations of abiraterone in the absence or presence of volasertib (7.5 nM), GSK461364 (5 nM), or onvansertib (15 nM). Relative viability was assessed and plotted as in (B). Similar to **Figure 2I**, multiple distinct Plk1 inhibitors synergize with abiraterone in C4-2 CRPC cells regardless of the presence or absence of androgens.

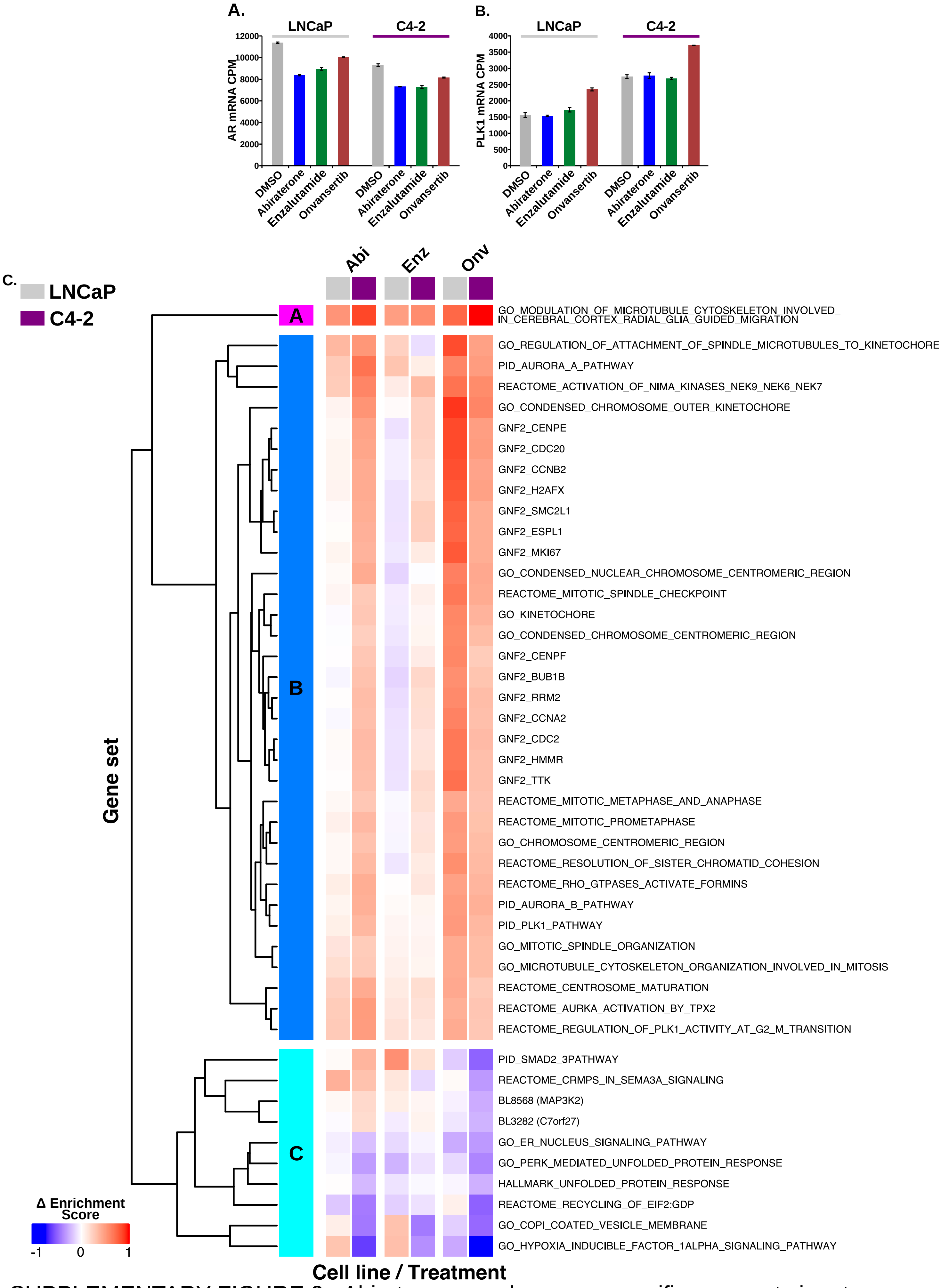


SUPPLEMENTARY FIGURE 2 - Abiraterone- and synergy-specific gene set signature identified from RNA seq analysis of prostate cancer cells.

**Supplemental Figure 2. Abiraterone- and synergy-specific gene set signature identified from RNA sequencing analysis of prostate cancer cells.**

**(A)** AR transcript levels measured in the RNA sequencing experiment described in **Figure 3** in C4-2 and LNCaP cells after treatment with the indicated drugs.

**(B)** Plk1 transcript levels measured in the RNA sequencing experiment described in **Figure 3** in C4-2 and LNCaP cells after treatment with the indicated drugs.

**(C)** Expanded and labeled version of the 45 gene set signature also shown in the **Figure 3F** heatmap. The expression of these gene sets were significantly altered by both abiraterone and onvansertib only in C4-2 cells and not in LNCaP cells. The majority of the gene sets refer to cellular components, biological processes and gene neighborhoods that are related to mitosis and mitotic spindle assembly. It is noteworthy that they are upregulated by abiraterone but not enzalutamide treatment in C4-2 cells, and neither antiandrogen increases their expression in LNCaP cells.

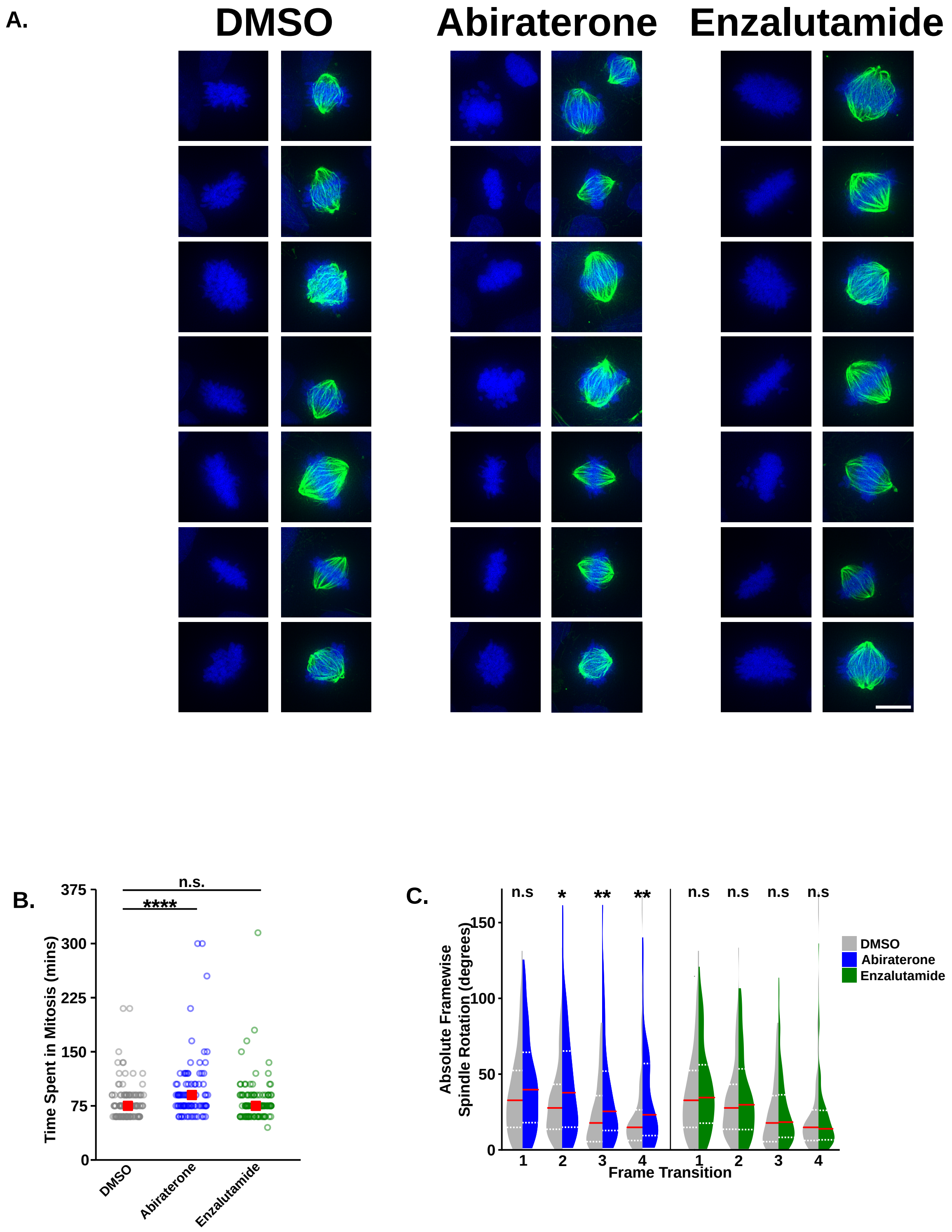


SUPPLEMENTARY FIGURE 3 - AR-independent effects of abiraterone on mitotic spindle assembly and chromatin condensation

**Supplemental Figure 3. AR-independent effects of abiraterone on mitotic spindle assembly and chromatin condensation.**

**(A)** Related to **Figure 4A**. Seven additional examples mitotic spindle morphology after treatment with DMSO, abiraterone, or enzalutamide for 16 hours. Fixed cells were stained with antibodies against tubulin (green) and DAPI for DNA visualization (blue). DMSO and enzalutamide treated cells had well-defined chromosome arms whereas abiraterone treated cells displayed ill-defined and apparently decondensed chromosome arms in mitotic cells. Scale bar lower right represents 10 μm.

**(B)** Related to **Figure 4E**. C4-2 cells expressing H2B-mCherry and mEmerald-tubulin were grown in media containing FBS, subjected to vehicle control, abiraterone (5 μM), or enzalutamide (10 μM), and then analyzed by time-lapse live-cell microscopy. The duration of mitosis for individual cells was then determined by image analysis ($n \geq 92$). **** $p \leq 0.0001$, n.s. not significant using a two-tailed Mann-Whitney U test.

**(C)** The distribution of absolute rotations between frames during mitotic progression, where frame transition one represents the amount of spindle rotation between when a mitotic spindle was first apparent to the second frame. On the left are violin plots comparing vehicle control to abiraterone, on the right is the same comparison with enzalutamide. Red line represents the median and dotted white lines are upper and lower quartiles. * $p \leq 0.05$, ** $p \leq 0.01$, n.s. not significant using a two-tailed Mann-Whitney U test.

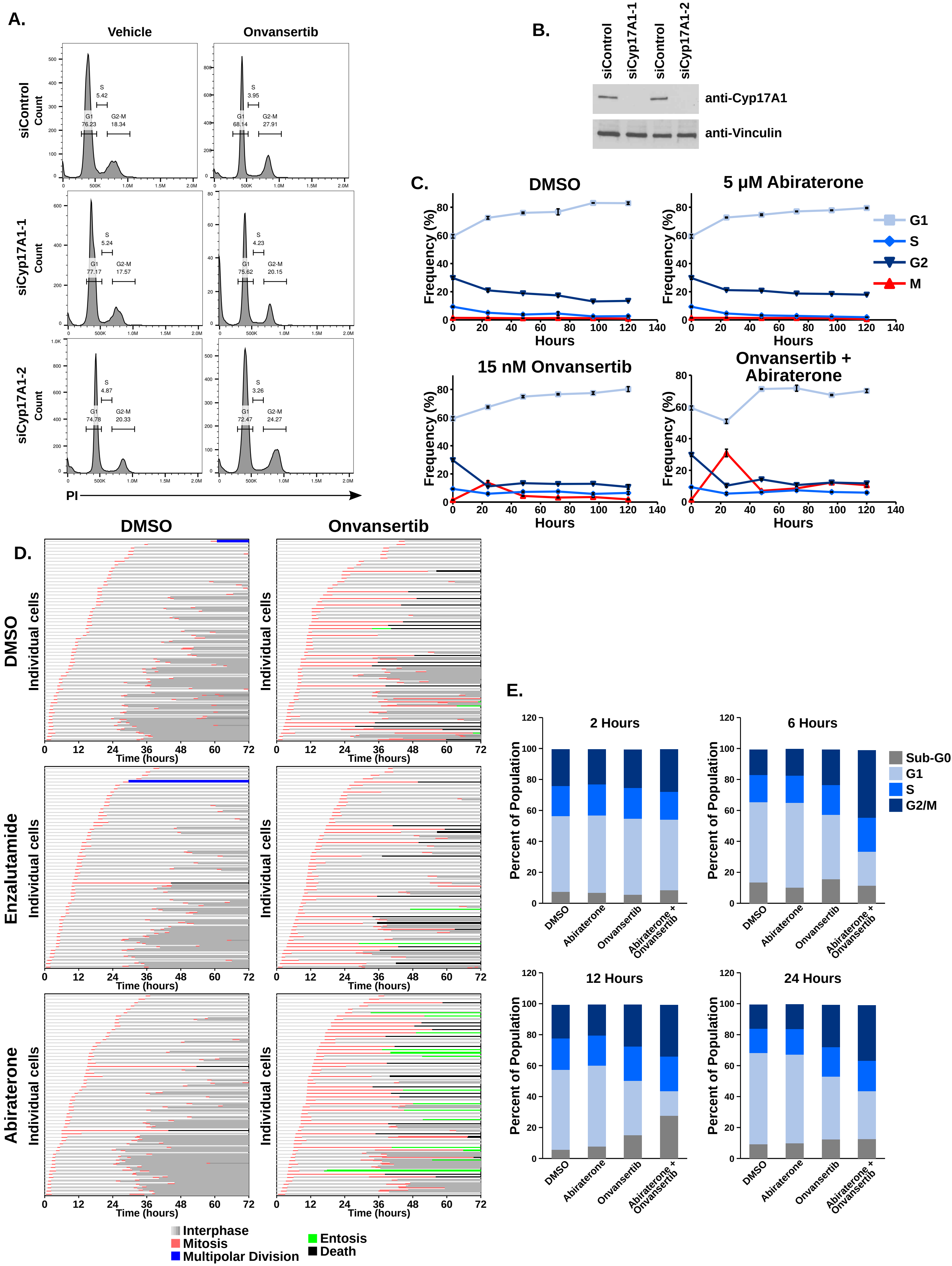


SUPPLEMENTARY FIGURE 4 - AR- and androgen-independent effects of abiraterone, onvansertib, and the combination on mitosis and overall cell cycle distribution in prostate cancer cells

**Supplemental Figure 4. AR- and androgen-independent effects of abiraterone, onvansertib, and the combination on mitosis and cell cycle distribution in prostate cancer cells.**

**(A)** C4-2 CRPC cells were transiently transfected with a control siRNA or one of two Cyp17A1-targeting siRNAs for 72 hours, and then subjected to 20 nM onvansertib for 6 hours prior to fixation. Cell cycle position was determined by flow cytometry using propidium iodide staining for DNA content. Shown are histograms depicting the distribution of DNA content. Gating was performed to separate G1, S and G2-M cells and their relative frequency among those gates is provided.

**(B)** Immunoblot confirmation of Cyp17A1 knockdown in C4-2 cells 72 hours after siRNA transfection.

**(C)** These data are related to **Figure 5C**, in which we analyzed the percentage of mitotic C4-2 CRPC cells over a time course after treatment with abiraterone, onvansertib, and the combination. Cells were collected, fixed stained with DAPI and antibodies raised against pHH3, and then analyzed by flow cytometry. Mean ± SEM (n = 3). In **Figure 5C**, we display only the percent mitotic. Here we provide percent G1, S, G2, and M phase cells over time in the four conditions.

**(D)** Related to **Figure 5D**, however, includes lines that depict duration of interphase of individual cells. Time-lapse microscopy of C4-2 cells expressing H2B-mCherry and mEmerald-tubulin treated with abiraterone (5 μM), enzalutamide (10 μM), and onvansertib (15 nM), and the combinations. For each condition, 60 cells were analyzed to determine both the duration of mitosis and the cellular phenotypes associated with mitotic arrest. Each row begins by tracking an individual cell in interphase (light grey), after completion of mitosis (red), the two daughter cells were tracked throughout the subsequent interphase (medium grey) which themselves could go through mitosis (red) to create up to four daughter cells (dark grey). Prolonged mitotic arrest was often followed by cell death (black) or entosis (green).

**(E)** DU145 AR-negative prostate cancer cells were treated with DMSO, 10 μM abiraterone, 10 nM onvansertib, or the combination for the indicated time prior to fixation. Cells were then stained with propidium iodide and their DNA content analyzed by flow cytometry as an indicator of cell cycle position.

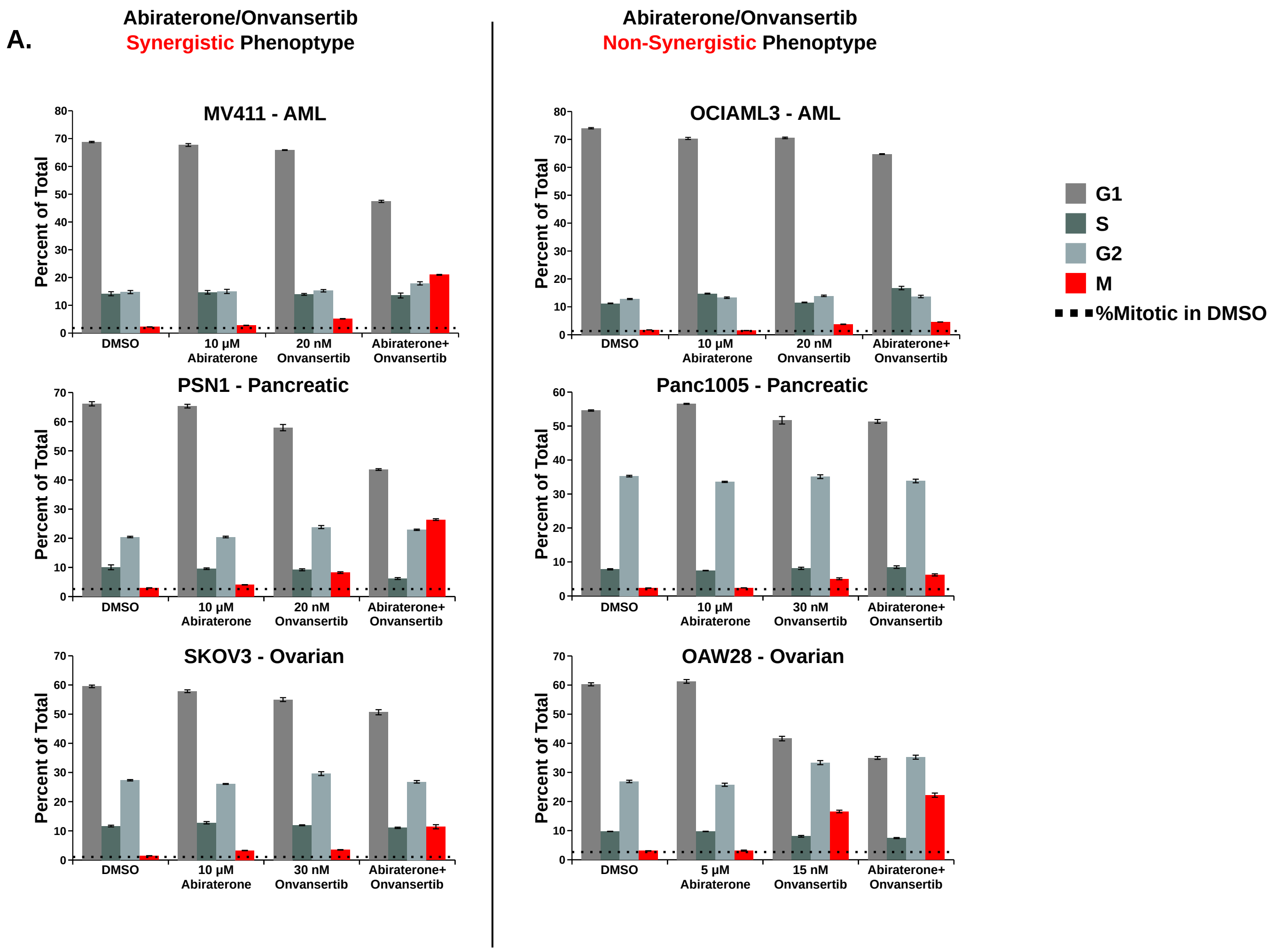


SUPPLEMENTARY FIGURE 5 - Effects of abiraterone, onvansertib, and the combination on AR-negative non-prostate cancer cells

**Supplemental Figure 5. Effects of abiraterone, onvansertib, and the combination on AR-negative non-prostate cancer cells.**

**(A)** These data are related to **Figure 6C**. The indicated cell lines were treated with the indicated drugs for 16 hours, fixed and then stained with DAPI and antibodies against pHH3 to assess cell-cycle distribution by flow cytometry. These graphs show the percentage of cells in each stage of the cell cycle in addition to the percentage of cells in mitosis presented in **Figure 6C**. Abiraterone treatment was 10 μM for all cell lines except OAW28, which was 5 μM. Onvansertib concentrations used were 20 nM for MV-4-11, OCI-AML-3, and PSN-1; 30 nM for Panc 10.05 and SK-OV-3; 15 nM for OAW28. Mean ± SEM (n = 3).

### SUPPLEMENTAL MATERIALS

#### Supplemental Movies

**Movie 1.** Chromatin structure in vehicle treated cells

**Movie 2.** Chromatin structure in abiraterone treated cells

**Movie 3.** Chromatin structure in enzalutamide treated cells

**Movie 4.** Identification of paired centromeres from sister chromatids

**Movie 5.** Video montage comparing mitosis of vehicle control with abiraterone treated cells

**Movie 6.** Normal mitotic cell division

**Movie 7.** Cellular division that occurred in a manner that was not parallel to the growth surface

**Movie 8.** Example of mitotic arrest and cell death caused by abiraterone and onvansertib

**Movie 9.** Entosis of mitotic cells
